# Environmentally dependent and independent control of cell shape determination by Rho GTPase regulators in melanoma

**DOI:** 10.1101/2021.10.11.463377

**Authors:** L. G. Dent, N. Curry, H. Sparks, V. Bousgouni, V. Maioli, S. Kumar, I. Munro, C. Dunsby, C. Bakal

## Abstract

In order to invade 3D tissues, cancer cells dynamically change cell morphology in response to geometric and mechanical cues in the environment. But how cells determine their shape in 3D versus 2D environments is poorly understood. Studying 2D versus 3D single cell shape determination has historically been technically difficult due to the lack of methodologies to directly compare the two environments. We developed an approach to study cell shape in 2D versus 3D by measuring cell shape at different depths in collagen using stage-scanning oblique plane microscopy (ssOPM). We find characteristic shape changes occur in melanoma cells depending on whether a cell is attached to a 2D surface or 3D environment, and that these changes can be modulated by Rho GTPase regulatory proteins. Our data suggest that regulation of cell protrusivity undergoes a ‘switch’ of control between different Rho GTPase regulators depending on the physical microenvironment.

## Introduction

The ability of metastatic cancer cells to invade three-dimensional (3D) structures such as tissues and organs is dependent on their ability to change shape in response to the environment. To respond to the environment, cells detect factors such as stiffness and geometry and in turn dynamically regulate their cytoskeleton [1–6]. Cells appear to convert between two major modes of migration and shape control: (i) adhesion based mesenchymal migration, (ii) cortical tension based ameboid and lobopodial migration [7–11].

On rigid 2D substrates such as glass (modulus of elasticity 60-64 GPa) [12], melanoma cells typically adopt an elongated ‘spindle’ or mesenchymal shape. In contrast, on softer environments such as collagen hydrogels, cells can adopt either mesenchymal or amoeboid shapes depending on environmental parameters such as stiffness and pore size [13]. The modulus of elasticity for different compositions of collagen hydrogel is a major determinant of cell shape and typically ranges between 50 and 5000 Pascals [14–19]. In stiff 3D collagen environments with small pore sizes that restrict migration, cells adopt mesenchymal modes of migration. Within soft or more porous 3D substrates cells tend to adopt ‘amoeboid’ and ‘lobopodial’ forms. Amoeboid cell migration is characterized by extensive contraction of cortical actomyosin and weaker adhesion to the substrate [20–22]. Amoeboid cells invade 3D matrices by pushing via ‘blebs’, which are protrusions in the plasma membrane generated by hydrostatic pressure. The ability of cells to switch between spindle and ameboid forms provides cancer cells with the ability to invade substrates with different stiffness and geometry [7,23].

Many of the changes in cell shape between environments of different stiffness and geometry are controlled by Rho GTPase proteins. For example, activation of RHOA is associated with increased myosin II-mediated contractility and cell rounding. In contrast, activation of CDC42 leads to WASP and arp2/3 activity and the formation of protrusions, whereas RAC1 activation leads to increased WAVE activity and the formation of protrusions in 3D spindle cells, or lamellipodia in 2D cells [24,25].

Although Rho GTPases are important determinants of cell shape, additional layers of regulation are necessary for GTPases to be able to ‘detect’ differences in the environment and respond with dynamic changes in activity. This fine-tuned regulation comes from Rho GTP exchange factors (RhoGEFs) and Rho GTPase activating proteins (RhoGAPs) [26]. RhoGEFs increase the activity of GTPases by promoting the release of GDP and loading of GTP. RhoGAPs (Rho GTPase activating proteins) decrease the activity of GTPases by catalysing hydrolysis of GTP to GDP [27]. Previously, RhoGEFs and RhoGAPs have been shown to be able to confer environmental responsiveness to Rho GTPases through recruitment and activation at distinct subcellular locations. For example, ARHGEF7 and SRGAP1 play important roles in regulating cell shape in 3D collagen versus fibronectin gels [28]. Differential activation of RhoGEFs and RhoGAPs likely underpin the ability of metastatic cells to change shape as cells transition between different environments such as tumor and normal tissue. Despite this, in many instances the RhoGEFS/GAPs that allow cancer cells to respond to a particular environmental context, remain to be identified.

Identifying the RhoGEFs, RhoGAPS that control different aspects of cell shape and migration in response to the environment is a major goal of biology, but there are significant challenges. For example, although disruption of the classical Rho GTPases such as RHO, RAC and CDC42 each have profound and stereotypic consequences for cell shape, the effect of RhoGEFs/GAPs can be more subtle and varied. In contrast to the ~20 mammalian Rho GTPases [29], there are some 145 Rho-regulatory GEFs and GAPs and their disruption often results in context specific modulation of shape.

A challenge in understanding the differences in shape control between 2D versus 3D environments is the difficulty of imaging the same cell populations in distinct environments simultaneously. In part this can be accomplished by culturing cells in invasion assays where cells migrate ‘up’ from a 2D environment into 3D collagen hydrogels [11]. However, conventional microscopy is poorly suited to imaging cells in both 2D and 3D environments, and is not appropriate for measuring 3D geometry.

3D imaging of cells at multiple depths can be achieved at high speed using light-sheet fluorescence microscopy (LSFM). Here we used stage-scanning oblique plane microscopy (ssOPM) for imaging melanoma cells invading a collagen hydrogel. ssOPM has been previously applied to time-lapse imaging of spheroids in multi-well plates [30]. This technique is based on oblique plane microscopy (OPM) where the same high NA objective delivers the light sheet and collects the fluorescence [31]. This system is built around a standard microscope frame and uses standard multiwell plates. We retain the advantages of working with a standard microscope frame but gain fast 3D imaging. The ssOPM images large volumes (4.2×0.32×0.144 mm^3^) corresponding to 100s of cells in 108 s. The collection NA is 0.7 and the system provides a spatial resolution of 0.5×0.5×5 μm^3^ [32]. This allows the 3D position and shape of large numbers of cells to be measured.

We analysed cell shape in control treated cells, and in response to depletion of different Rho-regulatory proteins. We found characteristic shape changes as cells transition from a 2D to 3D environment, and that these changes can be modulated by depletion of Rho-regulators. We also found that some Rho-regulators influence cell shape in a range of physical settings, while others are more context specific. In particular, focusing on cell protrusivity we found that depletion of Rho-regulators such as *TIAM2* changed protrusivity in both 2D and 3D environments, whereas depletion of *FARP1* only modulated protrusivity in 2D environments. This data suggests that cells can adjust shape control between different Rho-regulators depending on their local environment.

Taken together, these results reveal new context specific regulators of protrusivity and highlight the ability of high-throughput plate based volumetric imaging to rapidly assay and identify proteins in control of cell shape.

## Methods

### Cell culture

WM266.4 melanoma cells expressing CAAX-GFP (donated by the Marshall lab) were grown in Dulbecco’s Modified Eagle Medium (DMEM) supplemented with 10% heat-inactivated bovine serum (FBS) and 1% penicillin/streptomycin in T75 flasks. Cells were cultured at 37°C and supplemented with 5% CO_2_ in humidified incubators.

### Cell treatments and preparation

WM266.4 melanoma cells expressing CAAX-GFP were reverse transfected with OnTARGETplus SMARTpool (Table 1). Gene name and Dharmacon catalogue numbers were as follows: *ARHGEF35*, *ARHGEF9*, *PREX2*, *FARP1*, *TIAM2*, *SRGAP1*, *DOCK5*, *RND3* and *ECT2* (Dharmacon cat # L-032365-02-0005, # L-020314-00-0005, # L-014602-00-0005, # L-008519­00-0005, # L-008434-00-0005, # L-026974-00-0005, # L-018931-00-0005, # L-007794-00-0005 and # L-006450-00-0005) at stock concentration 20 µM in 6 well plates. Transfections were carried out using Lipofectamine RNAimax (Invitrogen) according to the manufacturer’s instructions. On the second day after transfection, 10^5^ cells/ml were re-suspended in 500 µl of 2.3 mgs/ml collagen rat tail (Gibco). A 100 µl volume of the collagen and cell mixture was dispensed in quadruplicate wells onto poly-D-Lysine (0.1mg/ml) coated glass bottom view 96 well plates (PerkinElmer). Plates were centrifuged @1200 rpm for 5 minutes at 4 ^°^C and incubated overnight in a tissue culture incubator. After incubation, cells were fixed with 4% PFA methanol free for 30 mins at RT. Wells were stained with DRAQ5 at a concentration of 5 µM, to label nuclei.

**Table 1.**
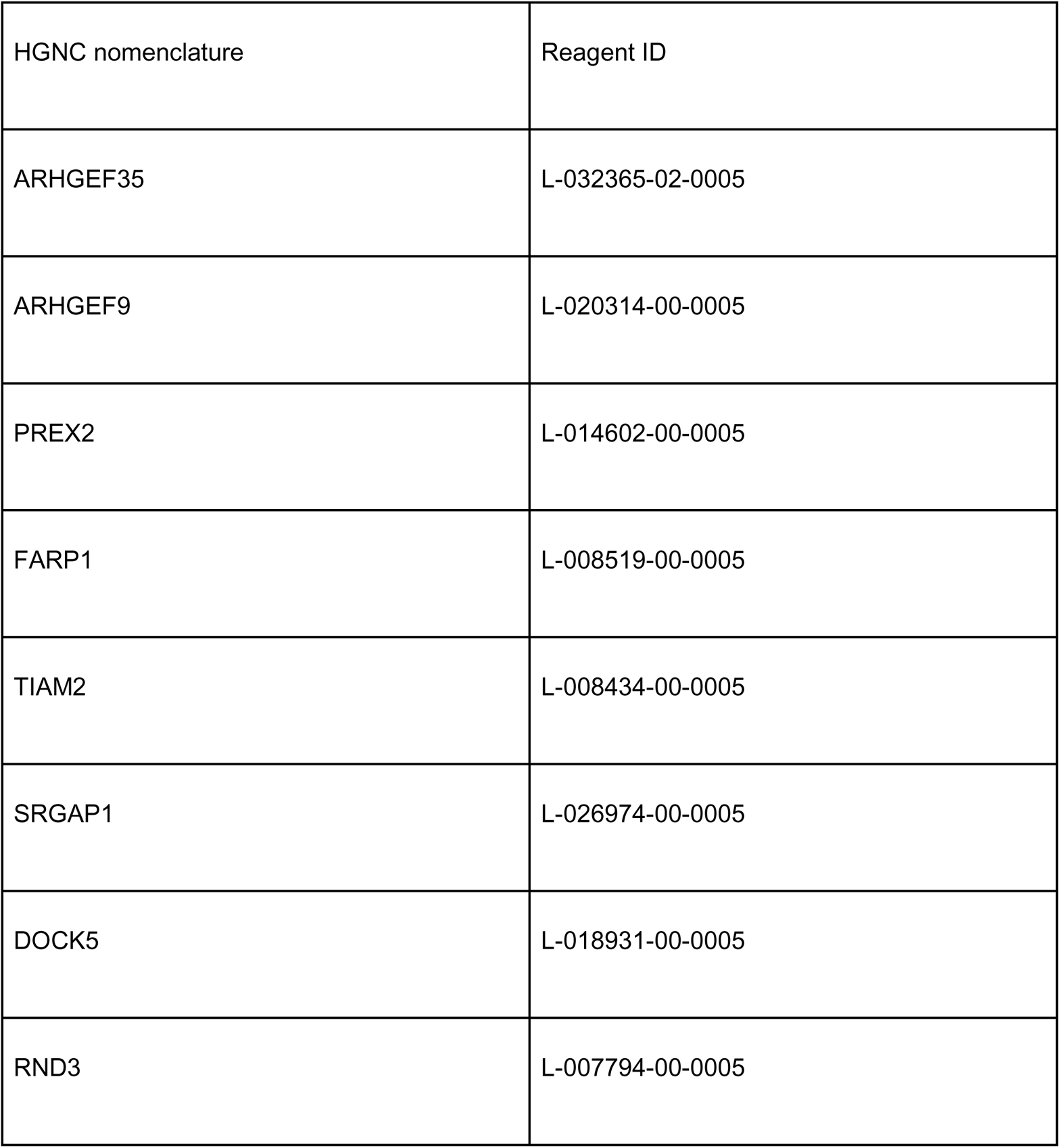
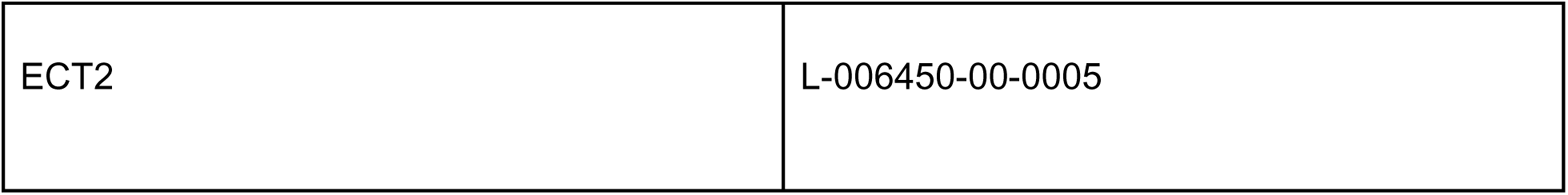
Gene names and siRNA reagent details for this study

### Microscopy setup

Oblique plane microscopy was performed using a setup reported previously [30] and a schematic is shown (Supplementary Fig 1A). Five excitation sources at different wavelengths (457, 488, 515, 561, 642 nm) are combined onto a common optical path using dichroic mirrors. An acousto-optic tunable filter – allowing switching and power control – is used to selectively couple the beams into a single-mode polarisation-maintaining fibre.

Light exiting the fibre is collimated (L1) then focused in the horizontal direction by a cylindrical lens (C1) onto the back focal plane of spherical lens L2. This results in a vertically orientated light sheet at 55 degrees to the optical axis of O2. The microscope formed by O2 and TL2 relays the light sheet to the image plane of the camera port of a commercially available microscope frame (Olympus IX71). The commercial microscope (comprising TL1 and O1) relays the light sheet to the image plane of O1. O1 is a 60X/1.2NA water immersion objective (Olympus UPLSAPO60XW). The microscope objective was fitted with a collar to provide a continuous supply of water immersion liquid.

The overall magnification between the image planes of O1 and O2 is set to be equal to the ratio of the refractive indices for the images formed at O1 (water) and O2 (air) to ensure that the lateral and axial magnification between the focal planes of O1 and O2 are equal [33]. The excitation light sheet produced across the focal plane of O1 excites fluorescence from the sample. The resulting fluorescence image is relayed back to the focal plane of O2. O3 is positioned at 35 degrees to the optical axes of O1 and O2, such that the fluorescence image of the region illuminated by the light sheet in the sample is perpendicular to its optical axis, and conjugate with the focal plane of O3. Together with tube lenses TL3a and TL3b, this system relays an image of the fluorescence emitted within the light sheet to the two sCMOS cameras. A dichroic beam splitter (DC) and emission filters (EM1, EM2) separate emitted fluorescence for two-colour imaging.

### Image acquisition

Stage-scanning OPM was implemented as described previously [34]. A motorised stage (SCAN-IM 120 × 80, Marzhäuser) controlled by a driver unit (Tango 2 fitted with AUX I/O option, Marzhäuser) outputs a TTL trigger each time it has travelled a predefined distance (1.4 μm for the results presented here). The TTL output is connected to a digital acquisition box (DAQ) (National Instruments NI USB-6229) configured to output a pattern of signals each time a TTL signal is received from the x-y stage. The signals control the laser power and illumination duration, and trigger the start of camera exposures. The stage scans in the y direction, as shown (Supplementary Fig 1A). During acquisition a scan speed of 0.1 μm ms^−1^ was used. A speed of 10 μm ms^−1^ was used to move between wells.

The two sCMOS cameras (PCO.edge, PCO) were both operated in Global Reset acquisition mode with 1280×1000 pixels. The exposure time was defined by the 2 ms laser exposure time. The two spectral channels were interleaved temporally to prevent cross-talk between channels. Per field of view, the x-y stage was scanned 4200 μm in the y direction (corresponding to 3000 frames per camera). Each field of view corresponded to 7.32 GB of data per channel and took 42 s to acquire. Image acquisition was controlled by a HP z840 PC with 128 GB of RAM and a 4×1 TB SSD configured in RAID 0. Saving and moving to the next well took a further 66 s (for a total of 108 seconds per well).

Volumetric imaging in 3 spectral channels (2 × fluorescence and 1 × scatter) was performed in two stages. The first stage acquired the two fluorescence spectral channels (CAAX-GFP and DRAQ5) for the entire plate. In the second stage, the image acquisition was repeated but now with scattered light from collagen imaged on camera 1 with 488 nm excitation and in the absence of an emission filter. Camera 2 was used to image the DRAQ5 channel for a second time. As images of DRAQ5 were acquired in both stages, they could then be used to measure any drift between image sets and thus enable the two sets to be co-registered.

### Image reslicing and registration

The average background level was measured for each camera by taking the average pixel value over a field of view acquired with no laser on. This was subtracted from the data prior to reslicing.

The 2D transform to co-register camera 1 and camera 2 was measured based on a 2-channel fluorescence image acquisition of a sample of 100 nm four-colour fluorescent beads (TetraSpeck, Thermofisher) in 10% agarose. The x-y shift, magnification and rotation needed to co-register the data acquired on camera 2 to that of camera 1 was measured manually using a custom script in MATLAB (imtranslate, imrotate, imscale and fliplr: Image Processing Toolbox, MATLAB). This transform was applied to all raw image data acquired on camera 2 prior to reslicing.

Raw ssOPM images are a set of image planes at 55 degrees to the optical axis of O1. The data was transformed into conventional coordinates (z parallel to the optical axis, x&y&z perpendicular). Reslicing was performed using a bi-linear resampling algorithm [35]. To increase speed, a custom-written Java implementation of the algorithm was used. For all image segmentation and cell shape analysis, raw camera images were binned by factor of 4 - prior to reslicing - to reduce data volume and analysis times. Images presented in figures in this paper were resliced with factor 2 binning. After reslicing voxel sizes were 1×1×1 μm^3^ (for analysis) and 0.5×0.5×0.5 μm^3^ (for display).

The collagen channel was coregistered in 3D with the fluorescence channels using the imregtform (Image Processing Toolbox, MATLAB) using the default optimizer parameters. The transform was measured on the DRAQ5 channel, which was common to both acquisitions, and applied to the collagen channel using the imwarp function in MATLAB.

### 3D rendering

3D renders are displayed as a 3D projection with trilinear interpolation using the Volume Viewer 2.01 Fiji plugin [36]. 3D surface renders were generated using the 3D viewer in Fiji.

### Image segmentation

Prior to segmentation, the collagen channel was viewed manually. Any volumes where the collagen was not present throughout the entire volume were rejected from analysis. Example (accepted) volumes are shown (Supplementary Fig 3A).

To verify the robustness of the 3D segmentation used in this paper, two methods were tested. An intensity-threshold-based approach using an Otsu threshold and an active contour method, which uses energy minimisation (Supplementary Fig 4A). Both methods generate a mask with minimal user input so can be applied to large datasets. For both methods, cells and nuclei were segmented in 3D.

Prior to segmentation, the tips of the parallelepiped-shaped volume imaged by the ssOPM image acquisition were removed by cropping in the y direction. This removed any parts of the volume which were not imaged over its full axial extent due to the light-sheet angle.

In the intensity-based method, thresholds were measured automatically for each field of view. The nucleus threshold was selected using Otsu’s method with a single level (multithresh, Image Processing Toolbox, MATLAB). The cell body was masked using a similar method. In this case there were 3 intensity levels in the image, background, brightly fluorescent cell membrane and dim fluorescent protrusions. To include all parts of the cell in the final mask, the lowest threshold found by a two-level Otsu method was used.

In the active contour method, an initial guess of the mask was generated using a threshold of 5 digital numbers (just above the background). The final mask was formed after 500 iterations (nuclei) or 1000 iterations (cell) of the active contour method (Image Processing Toolbox, MATLAB).

The final nucleus mask was generated for both methods by separating touching nuclei. To achieve this, the Euclidean distance transform (bwdist, Image Processing Toolbox, MATLAB) was used on the inverse of the mask to determine the distance of each voxel to the edge of the 3D mask. A watershed (Image Processing Toolbox, MATLAB) was used on the negative of the distance transformed image to separate nuclei based on regions where the mask narrows. Nuclei with volume less than 250 μm^3^ were rejected at this stage.

As the nuclei are part of the cell, an OR operation is applied to the cell mask and the nucleus mask to generate a combined mask. Any connected components in the cell binary mask which do not contain a nucleus were rejected. Touching cells are separated using a marker-based watershed approach. Nuclei are set as the low points (digital value 0), the cell body (digital value 1) as intermediate points and the background as high points (digital value infinity). The watershed finds the halfway point between touching nuclei. The watershed then underwent an AND operation with the original cell mask to generate a final mask. Following segmentation, cells touching the edges of the image volume are removed. Cells with volume below 512 μm^3^ were rejected. Nuclei of rejected cells were removed from the nucleus mask.

### Image measures

Cell and nucleus shape measures were read out using the regionprops3 (Image Processing Toolbox, MATLAB). Further statistics were derived from the outputs of this function as described in Supplementary Fig 2A.

### Data processing

#### Outlier removal

Imaging of 3 plates produced a segmented dataset of more than 3×10^4^ cells. We aimed to remove cells and nuclei that could not be segmented accurately due to low expression of the CAAX-GFP transgene, as well as cells with surface areas that are conspicuously large due to under-segmentation. To remove cells that were improperly segmented due to low expression of the CAAX-GFP construct, we removed cells that had both: (i) low CAAX-GFP intensity (an average intensity of less than 1000 in camera digital numbers); and (ii) a nuclear to total cell volume ratio of greater than 0.95. These cells were removed from subsequent analysis as they represent nuclei with cells that cannot be appropriately measured due to low CAAX-GFP transgene expression. Applying these criteria removed approximately 500 cells (~ 1.8 % of original total).

#### Coverslip localisation

The position of the coverslip was estimated using the fluorescence in the nucleus channel. To account for spatial variations in the axial position of the coverslip over the field of view, the nucleus channel was divided into 16 segments in the y direction and the average signal in each x-y plane was found for each z position in each segment. For each segment, the coverslip location was defined as the point when the signal first reaches 45% of its maximum value when moving in the positive z direction. A full 2D map of coverslip height was then produced from the 16 measurements using bilinear interpolation, giving a smooth change of coverslip height across the field of view. This map of coverslip height reflects the spatial variation across the field of view. However, this is a relatively crude method and does not account for variations in nuclear intensity between knockdowns and between plates, therefore there is a remaining global offset in coverslip position for each well that is accounted for in the next step.

#### Nucleus height determination

Following coverslip position estimation, we calculated the lower boundary of the mask for each nucleus above the coverslip (the bottom of the nucleus). To remove the remaining global offset in coverslip position, we found the nucleus with the lowest calculated height in each well and subtracted this value from the height of every cell in that well so that the lowest cell in each well had a height of zero. Finally, we corrected for wells where microscopic detachment of collagen (less than 6 microns) from the well bottom had occurred after fixation but before imaging. These wells were identifiable by discontinuities in nuclear height distributions between cells at a lowest position (attached to coverslip) and a larger number of cells in the lowest extent of the collagen gel. In these cases the lowest positioned cell in the collagen was also registered to the height of the coverslip. The median position adjustment was 1.17 micrometers.

#### Feature reduction and feature normalisation

We originally computed more than 20 measurements of cell and nuclear shape features, and subsequently reduced this set to a set of four features. We used clustering to remove the most highly correlated features as follows. First, we used single cell data to calculate Pearson correlation values between each feature using the ‘cor()’ function in R. Pearson correlation values between features were hierarchically clustered using the hclust() function in R, and the ‘complete’ linkage method (see Fig 2). The resulting cluster was partitioned into four groups, and a single cell or nuclear feature was chosen as a representative feature for each group. To allow for comparison between plates that were prepared and imaged on different days, we normalised single cell measurements for cell and nuclear features within each plate. We performed normalisation by dividing feature measurements for a single cell, by the plate median across all conditions for cells in that feature.

**Fig 2.**
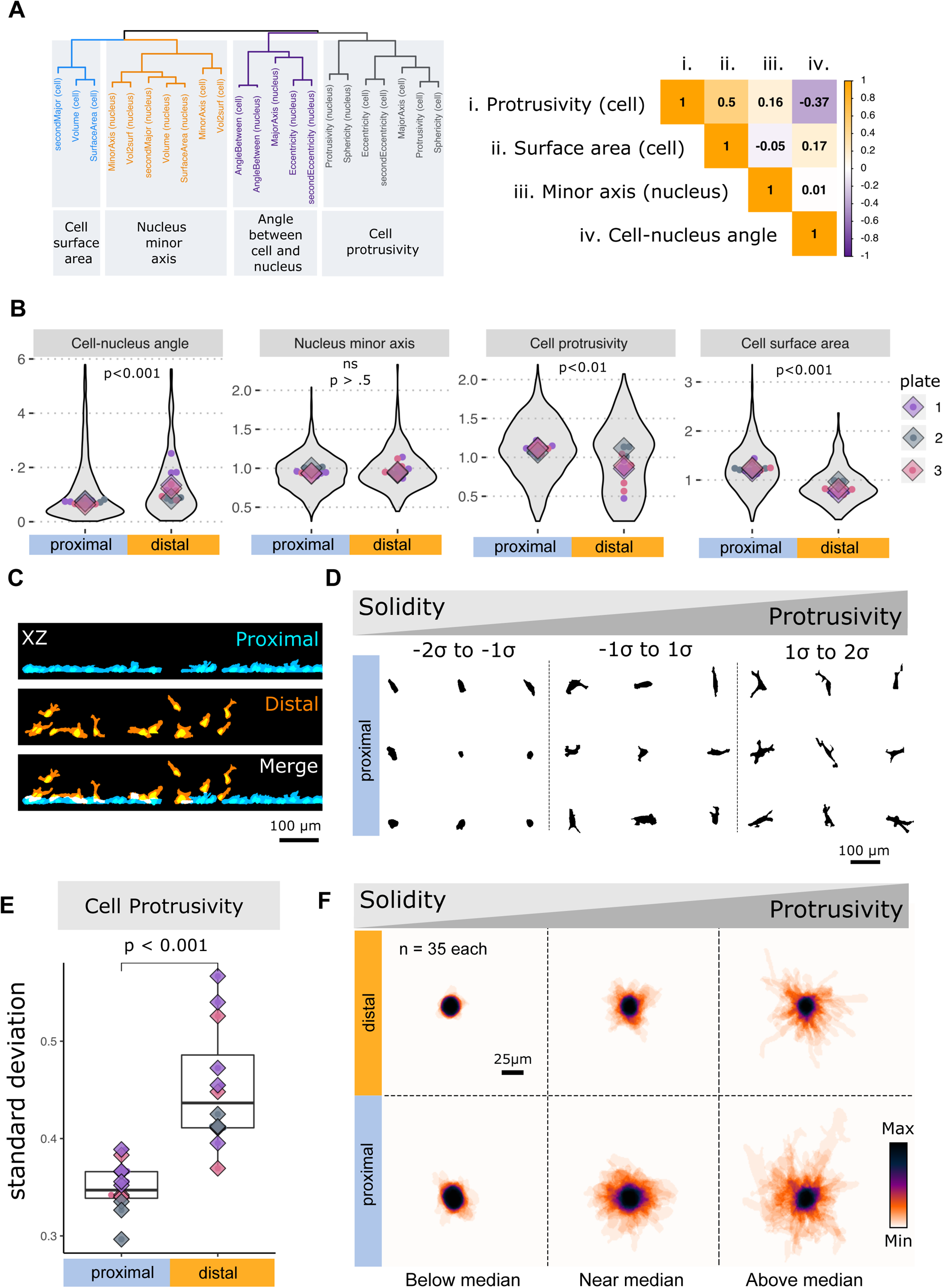
There are characteristic shape changes between cells with nuclei positioned away from a rigid substrate. **A.** Hierarchical clustering of cell and nuclear features, showing partitioning into four groups of correlated features (top row of grey boxes), and the feature selected to represent each group (bottom row of grey boxes). Right hand side shows a correlation matrix between the four shape features selected for analysis (also labelled ‘i’,’ii’,’iii’ and ‘iv’). The matrix indicates the direction and magnitude of correlation between these features. **B.** Comparison of shape features between cells with nuclei ‘proximal’ to the coverslip (blue) and ‘distal’ to the coverslip (orange). Cells with the base of nucleus less than 7 microns from the coverslip are proximal, and cells with the base of nuclei greater than 7 microns are distal. Violin plots show the distribution of single cells (grey filled region). Also shown are well medians (coloured points), and plate medians (coloured rhomboids). Each colour indicates data from a different experimental plate. Statistical tests are paired Wilcox tests of data from 3 independent experiments aggregated at the ‘well’ level. **C.** Cell segmentation masks from XZ MIPs from an exemplar field of view showing separation of cells into proximal and distal groups on the basis of nucleus distance from the coverslip. **D.** Examples of the heterogeneity in protrusivity that is inherent amongst proximal control treated cells. Plots of shape outlines from cells in three groups: below average (−2σ to −1σ); near average (−1σ to 1σ); and above average (1σ to 2σ). **E.** Comparison of the estimated probability density for protrusivity between proximal (blue shading) and distal cells (orange shading). Points show data at the well level. Point colours indicate replicate experimental plates, and are the same as in B. **F.** Images are stacked maximum intensity projections of cell outlines (as in D), for cells with protrusivity at different ranges from the median. n = 35 cells per projection.

### Data analysis

As some of the cell and nuclear shape features we examined did not have a normal distribution, we used non-parametric tests throughout this study. To test for differences in measures of central tendency between two conditions we used a paired Wilcoxon test with BH adjustment. For tests of difference in central tendency between more than two conditions we used a Kruskal-Wallis test, followed by a Dunn’s test to discern which conditions differed from control. Tests for differences between groups were performed on summary statistics calculated for wells, with multiple wells from three different plates in each test. Tests at the well level used at least 12 wells per treatment, from three plates, with at least 50 cells in the dataset per condition. Statistical tests were performed with functions in the R programming language and environment. Principal component analysis was performed on cell measurements aggregated at the well level, using the ‘prcomp()’ function in R.

### Confirmation that shape changes between environments are robust to segmentation method and light sheet PSF shape

#### Segmentation method

There has been an acceleration in recent years in the development of fast 3D microscopy techniques for imaging isolated cells. However there is limited software for segmentation of full 3D volumes. We test two methods for unsupervised segmentation of a full 3D dataset of cells. We tested, on control cells, a method which segments based on an Otsu threshold set for each field of view and an active contour method (both segmentation modes are described in detail in the Methods section). Supplementary Fig 4A shows the features measured using both segmentation methods. They are separated into coverslip proximal and distal groups based upon nucleus position with respect to the coverslip (as outlined in the previous section). For most of the features the Otsu and active contours methods give similar values. Changes between coverslip proximal and distal cells appeared consistent between threshold and active contour based segmentation. Overall this suggests that results are repeatable between segmentation methods. As either segmentation method is viable for this dataset, the Otsu threshold approach was chosen due to the shorter computation time.

#### Anisotropic PSF shape

In light-sheet microscopy, the point spread function (PSF) is usually not spherical. The FWHM spatial resolution of this ssOPM system has been previously reported as 0.5 um in the plane of the light sheet, with a light sheet thickness of 3.8 um at the waist. The light thickness increases to 5.4 μm over a distance of 50 μm from the centre of the field of view [37]. We set out to establish whether the anisotropic PSF might affect coverslip proximal cells - that are more likely to be flatter - differently to coverslip distal cells. We therefore eroded the cell and nucleus masks from all segmented data by an object approximating a worst-case PSF. This was performed using MATLAB’s imerode function with a morphological structuring element consisting of a 1×1×5 pixel kernel angled at 45° to the coverslip plane (closest possible approximation to a 55° light sheet angle). The voxel size in the image data was 1 um^3^, so this corresponds to a 1×1×5 um^3^ PSF.

Supplementary Fig 4B shows pair plots of cell features for the thresholded mask (normal) and the eroded (imerode) version. Coverslip proximal and distal cells have similar normalised distributions for normal and eroded masks. This suggests that, for the selected features, the anisotropy of the mask has limited impact on the comparison between coverslip proximal and distal cells.

#### Spatially varying light-sheet thickness

The ssOPM microscope uses an illumination light sheet with a confocal parameter of 100 μm. Therefore, the thickness of the light sheet varies as a function of the z range and is thinner at the centre of the z-range than the edges. Coverslip distal cells tend to be closer to the centre of the z range than coverslip proximal cells. Therefore cells on the coverslip experience a different PSF, which may affect the features measured.

To quantify this effect a reference sample of cells plated on the coverslip (no collagen) was used (referred to as the test plate). The same cells were imaged at three different z positions corresponding to the top, middle and bottom of the axial range. Example images are shown in supplementary Fig 5B. Cells from the test plate and collagen assay were grouped into bins (bottom, middle and top) based on their z coordinate (0-48, 48-96, 96-144 μm). Values were then normalised to the mean value for that feature in the lowest (first) bin.

Supplementary Fig 5A shows side by side plots of the features from the test plate and control cells from the collagen plates. Feature values were normalised by dividing by the average value in the bottom bin. On the test plate, cell surface area and the angle between cell and nucleus are both within one standard deviation of one at all heights, suggesting they are not significantly changed by the spatially varying PSF. Cell protrusivity increases in the middle bin of the test plate but a decrease in protrusivity is observed in the same bin on the collagen plate. This suggests that the decrease in protrusivity observed in coverslip distal cells is due to the change in physical environment and not due to the light sheet thickness. Nucleus minor axis shows a decrease below 1 in both the middle and top bins, however this decrease is not seen in the collagen plate. Cell and nucleus volume were also tested for the test plate, but this metric was found to be affected by the spatially varying light sheet thickness. This is expected for the thin flat cells used in the test plate, which represent a deliberate worst case, as these cells are generally thinner than the light-sheet thickness. The cell volume metric was therefore not used in the analysis in this paper. We concluded that the behaviours of ‘cell surface area’, ‘angle between’, ‘protrusivity’ and ‘nucleus minor axis’ in the collagen plate are not explainable by the PSF shape alone and are dominated by other factors.

## Results

### An experimental paradigm to measure cell shape in distinct physical environments

To study the differences in shape as cells transition between two mechanically and geometrically distinct environments, we suspended cells in 2 mg/ml (initial concentration) type 1 collagen (rat tail), seeded this mixture into glass-bottomed 96 well plates, and centrifuged them. We incubated cells for 24 hours before fixation. Glass is a rigid or ‘hard’ substrate (on the order of 1 GPa), whereas collagen at a concentration of 2 mg/ml is relatively elastic [15] (between 300 and 1600 Pascals) [38–42] (Fig 1A). We used this system to study control treated cells, and also cells treated with siRNA targeting a variety of Rho-regulators that we had selected from preliminary screening (Fig 1B). To compare cell response between physical environments we used stage scanning oblique plane microscopy (ssOPM) to image the geometry of cells with nuclei at different distances from the glass coverslip (Fig 1C-E). In each single volume, approximately 200 cells were imaged across a 144 μm z range. Using this imaging approach for all of the wells and treatments in our study generated an initial dataset containing more than 30,000 individual cells. A technical advantage of this system is that variables are internally controlled because comparisons can be made between 2D and 3D microenvironments for a large number of cells in the very same well of a 96 well plate. In the case of a single gene knockdown, protein depletion, media conditions, collagen concentration and biological composition between ‘proximal embedded’ and ‘distal embedded’ cells are shared between 2D and 3D cell microenvironments.

**Fig 1.**
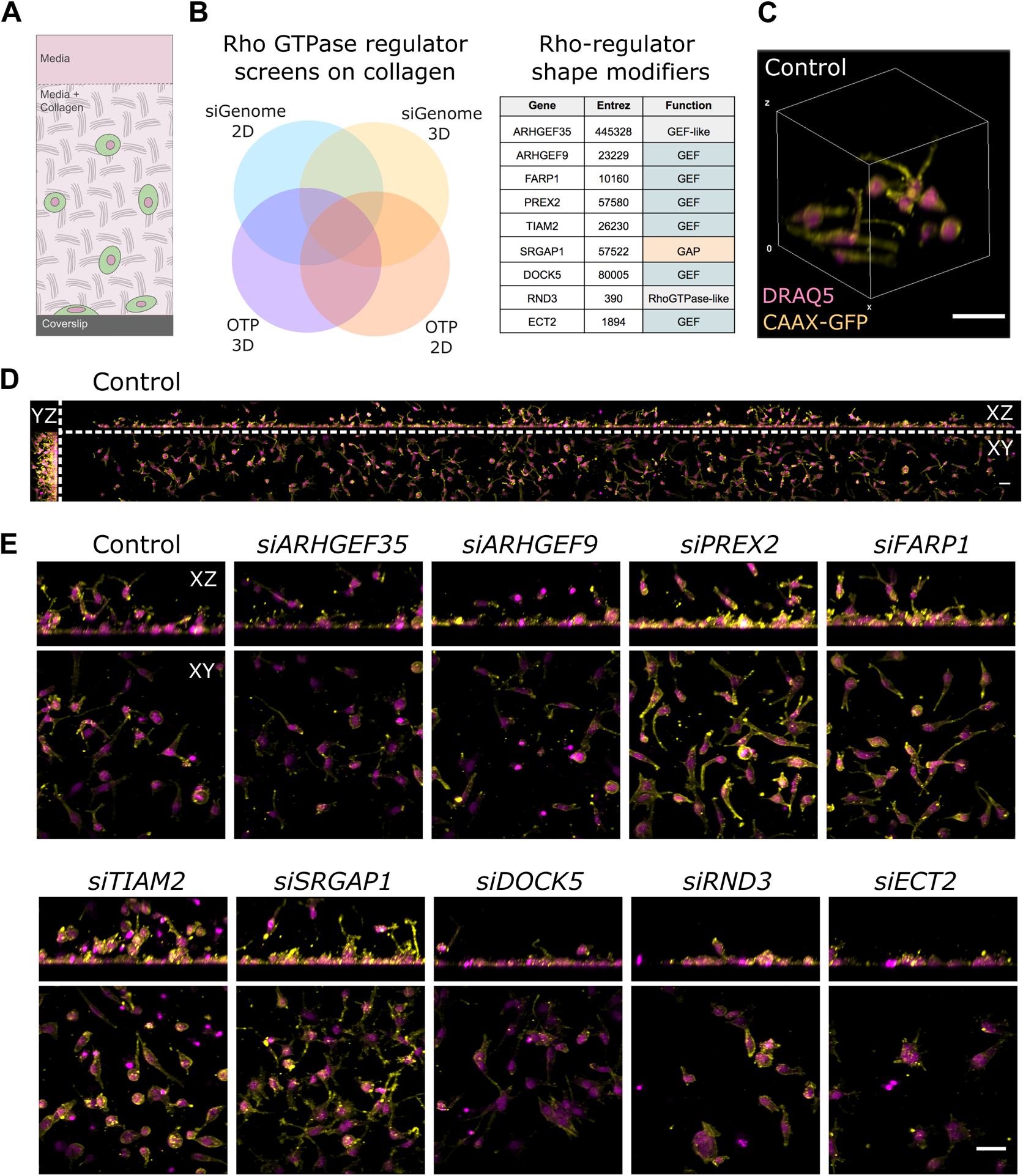
Genetic depletion of Rho-regulators in distinct physical contexts and imaging in 3D by oblique plane microscopy. **A.** Schematic illustration of WM266.4 melanoma cells embedded in collagen at different distances from coverslip. **B.** Schematic of preliminary shape screens on collagen and list of final Rho-regulators targeted for depletion. **C.** 3D volume rendering of control cells imaged by ssOPM. **D.** Maximum intensity projection (MIP) of the full field of view imaged for a single well. **E.** Example MIPs of zoomed regions for the nine different Rho-regulator siRNA treatments. Cells are marked by CAAX-GFP (yellow) and nuclei are marked by DRAQ5 (magenta). Images are from plate 2 row E. Scale bars are 100 microns.

To quantify cell morphology, we imaged GFP signal in CAAX-GFP-expressing WM266.4 cells, and visualized nuclei using DRAQ5 (Methods). Initially we generated over 20 measures of cell or nuclear shape features (Supplementary Fig 2A). We performed dimensionality reduction by using hierarchical clustering to group these measurements into four clusters of highly correlated shape features (Fig 2A). A representative shape feature was selected from each cluster to be used throughout our analysis. By considering the shape features in each cluster we were also able to suggest an interpretation of the underlying biology that is tracked by each feature (Fig 2A). The shape features chosen were ‘Cell surface area’, ‘Angle between cell and nucleus’, ‘Nucleus minor axis’, and ‘Cell protrusivity’ (description of features in Supplementary Fig 2A). After feature reduction, no features had an absolute correlation of greater than 0.5 (Fig 2A). Collectively we refer to these shape features as global geometry features.

### Human melanoma cells adopt different shapes in distinct microenvironments

To understand how cell shapes change based on distance from the glass coverslip we first examined untreated WM266.4 melanoma cells. The data analysed included more than 2,500 untreated melanoma cells in 12 wells across three plates. Based on cell shape measurements in our dataset, and on previous studies of hydrogels plated on stiff substrates we classified cells as ‘proximal’ to the coverslip when the base of the nucleus was less than 7 microns, and ‘distal’ if the base of the nucleus was greater than 7 microns from the coverslip. To search for changes in shape we compared global geometry features of proximal and distal cells (Fig 2B). This revealed significant stereotypic differences in cell morphology depending on whether cell nuclei were proximal or distal to the coverslip. These changes included reduced cell protrusivity, and smaller cell surface area in distal versus proximal cells (Fig 2B). In contrast to these cell shape features, the nuclear geometry feature, length of the minor axis of the nucleus was not different between proximal and distal cells (Fig 2B). Thus cells invading 3D collagen gels are typically less protrusive and have a smaller surface area than in 2D environments.

Although our nuclear shape measures were not altered, we found changes in the relationship of the cell to the nucleus. When cells were proximal to the coverslip the major axis of elongation of the cell and nucleus was coordinated or ‘coupled’. In contrast, for cells positioned away from the glass coverslip we found an increase in the angle between the cell and nucleus. This suggests the orientation of the nucleus was less constrained by cell geometry when positioned away from the glass coverslip.

Previous studies have noted that WM266.4 cells exhibit extensive heterogeneity in morphology when cultured either on stiff 2D plastic, or soft collagen matrices - adopting either amoeboid or spindle forms [8,43,44]. But how the extent of this variability changes between 2D and 3D is poorly understood. We tested whether variance in a range of shape features is changed between proximal and distal settings. Due to the connection between cell protrusivity and metastatic potential, we focused on variation in protrusivity (Fig 2E). To visualise these variations we grouped the cells into three groups - below average, near average and above average protrusivity - and selected 9 representative masks from the cell masks (Fig 2D). We also made a visual summary of variation in protrusivity (without grouping the cells) by creating ‘stacked maximum intensity projections’ (stacked-MIPs) (Fig 2F). The projected cells were the first ‘n’ cells in our dataset, which matched the protrusivity criteria. These stacked-MIPs support the finding that protrusivity is decreased when cells are positioned away from the coverslip. These observations are consistent with those made by ourselves and others that WM266.4 cells alternate between round and spindle forms in 3D gels, but are more homogenous on 2D stiff surfaces [7,8,44]. This increased variance may reflect that there are more degrees of freedom away from the rigid coverslip, which may make it less likely for cells to adopt stereotypic shapes.

### Rho-regulators disrupt differences in shape between proximal and distal cells

Having identified characteristic shape differences between cells that are proximal or distal to the coverslip, we interrogated the molecular control of these shape changes. To do this we depleted a range of Rho-regulatory proteins, and examined whether their function was affected by distance from the coverslip. To select a set of Rho-regulators for study by ssOPM, we had conducted a collection of preliminary screens on cells plated on collagen to look for genes that control cell morphology and imaged by confocal microscopy (Fig 1B). We identified nine Rho-regulators that have a potent influence on cell shape (Fig 1B).

We depleted these nine Rho-regulators and for eight of these we measured the effect on shape transitions between ‘proximal’ and ‘distal’ cells (Fig 1E and 3A and 3B). *ECT2* depleted cells frequently had a multinucleate phenotype consistent with failed cytokinesis (Fig 1E). This phenotype indicated potent protein depletion from our treatments, however to focus on primary effects of gene knockdown on shape we did not analyse these cells. To see how Rho-regulators contribute to cell shape in proximal 2D and distal 3D cells, we projected our four global geometry features (Fig 3A) in principal component (PC) space (Fig 3B). The two largest contributions to PC1 were cell protrusivity and cell surface area, while the two largest contributions to PC2 were the nucleus minor axis, and the angle between the cell and nucleus.

**Fig 3.**
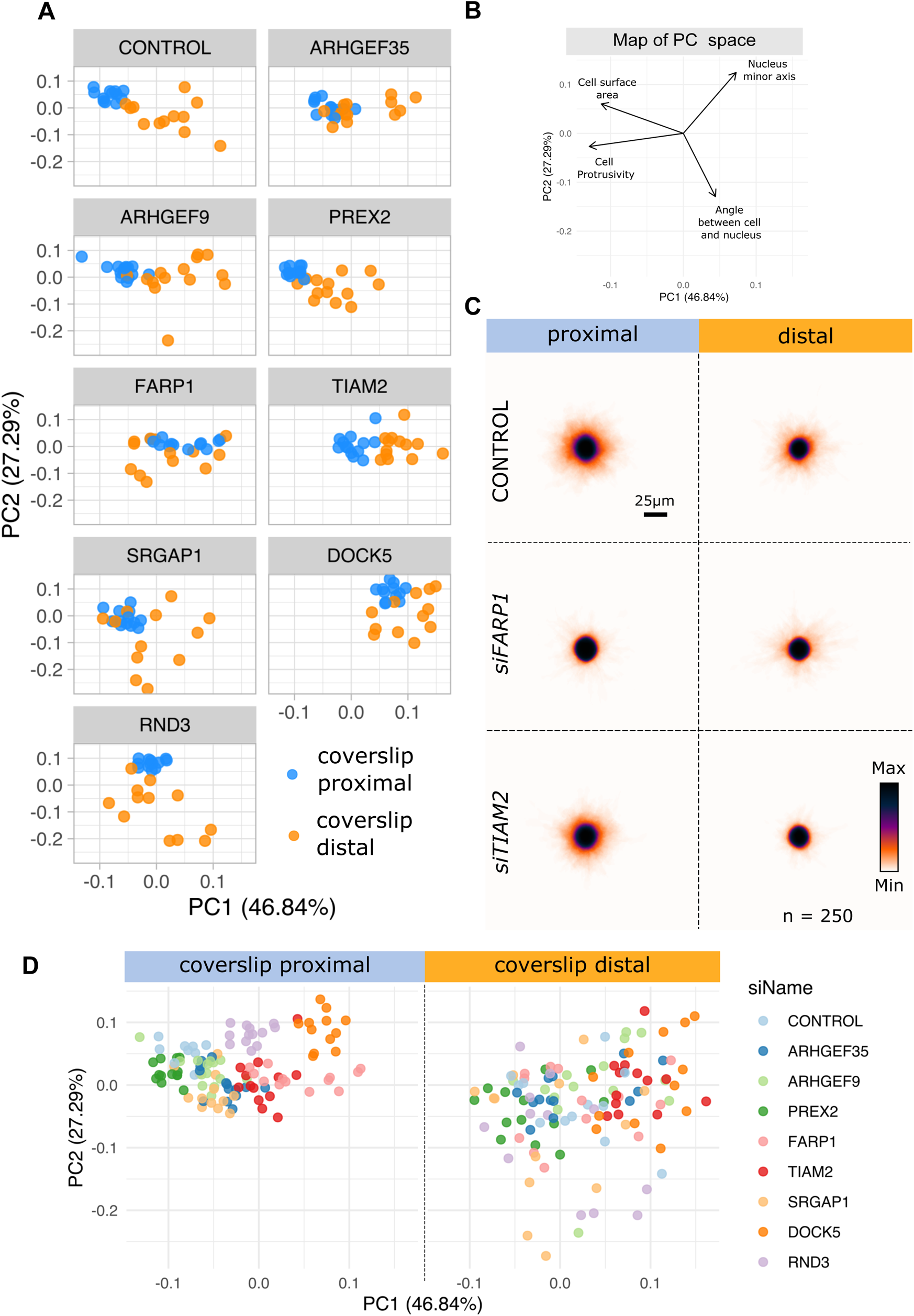
Depletion of Rho-regulators can disrupt shape differences between cells with nuclei at different distances from a rigid substrate. **A.** Principal components analysis (PCA) plotting of well median values for cell shape features showing proximal (blue) and distal (orange) cells. In *control* cells, proximal and distal cells are separated in PCA space. In contrast, treatments such as *FARP1* depletion result in an overlap between proximal and distal cells in PCA shape space. **B.** Map of how shape features project to PCA space. **C.** Stacked projections of cell segmentation masks for coverslip proximal or distal cells that are Control treated or depleted for *FARP1* or *TIAM2*. **D.** PCA plots comparing the effect of Rho-regulator depletion on shape in cells with nuclei proximal or distal to the coverslip.

For control cells in PC space, coverslip proximal cells formed a cluster with lower variance than coverslip distal cells. Coverslip distal cells explore shapes characterised by reduced protrusivity and cell surface area (higher PC1) and lower nucleus minor axis and greater angle between cell and nucleus (lower PC2). We found that depletion of some Rho-regulators generated overlap between proximal and distal cells in PC space (Fig 3A). For example, this overlap was substantial for cells depleted for *FARP1.* This was true but to a lesser extent for *ARHGEF35* and *SRGAP1­*depleted cells. The shape convergence we saw in PC space was visually supported by generating stacked-MIPs of 200 cells (Fig 3C), which indicated greater similarity between proximal and distal cells for *FARP1*, compared to control. Taken together we identify *FARP1*, *ARHGEF35* and *SRGAP1* as important for reducing differences in shape between proximal and distal contexts (Fig 3A-C).

Here we have shown that proximal and distal positioned WM266.4 melanoma cells are separable in PC space based on a small set of shape features, and this separation can be disrupted by depletion of some Rho-regulators.The ability to disrupt shape changes indicates that the shape differences between proximal and distal cells are not just biophysical responses to changes in the physical environment but are active transitions mediated (at least in part) by the signalling of Rho-regulators.

### The effect of Rho-regulators on cell shape is environmentally constrained

Visualisation in PC space also suggested that Rho-regulators have stronger and more stereotyped effects on cell shape in proximal compared to distal cells. In the proximal cells we found that our four global shape features were sufficient to separate many Rho-regulator depleted cells from control treated WM266.4 melanoma. This was especially the case for *SRGAP1*, *FARP1*, *TIAM2, RND3* and *DOCK5* (Fig 3D), where depletion of these proteins in proximal cells created combinations of shape features that were separable from control treated WM266.4 melanoma cells. This was in contrast to the distal context, where cell shape features overlapped between control WM266.4 melanoma and Rho-regulator depletion (Fig 3D), and where cell shapes were more widely distributed in PC space (Fig 3D).

This supported our finding that Rho-regulators have a potent and stereotypic effect on cell shape, but this effect depends on distance of the cell from the glass coverslip. Our data suggests that for distal cells the physical environment has an overarching control on shape and increases variability in cell shape.

### Control of cell protrusivity

We next sought to identify specific shape differences that are disrupted by Rho-regulator depletion, and focused on cell protrusivity. It is critical to understand regulation of cell protrusivity in WM266.4 melanoma because changes in protrusivity are linked to malignant cell migration. In the past a comprehensive understanding of the genetic control of cell protrusivity has been confounded by different control of protrusivity between 2D and 3D environments, as well as between rigid and soft environments. We define protrusivity as 1-ratio of the mask volume to its convex hull volume (Fig 4A & B). This is similar to the spreading metric used by Isogai *et al* [45]. Increases in our protrusivity metric correlate with the number of cell protrusions, but also the increased length of cell protrusions, and the angle between cell protrusions.

**Fig 4.**
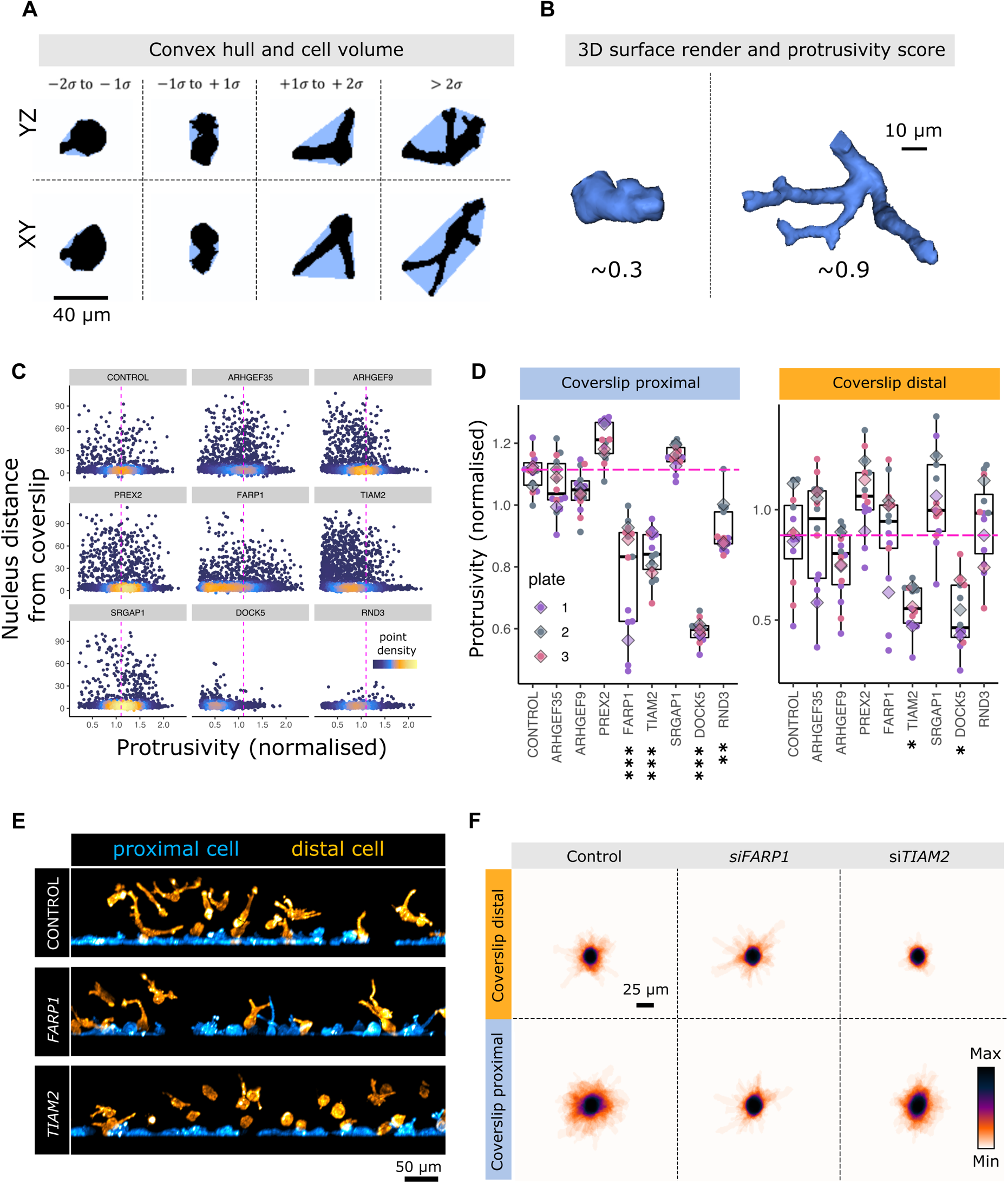
Depletion of Rho-regulators reveals genes that control protrusivity in broad and specific contexts. **A.** 2D Projections of cell segmentation masks (black) and convex hull (blue) used to calculate protrusivity. XY and YZ views are shown. **B.** 3D surface rendering of cells with different measures of protrusivity. **C.** Single cell plots of cell protrusivity against nucleus distance from the coverslip in control cells and cells depleted for Rho-regulators. Heatmap indicates density of points for overplotted regions of each chart. Horizontal dashed magenta line indicates median protrusivity across the entire dataset. **D.** Comparison of cell protrusivity between cells with depletion of a range of Rho-regulators for cells with nuclei proximal to the coverslip or distal to the coverslip. Data are aggregated and plotted at the well level (boxplots and scatterplot points), and at the plate level (rhomboids) The different colours indicate data from different plates. Statistical tests are Kruskal-Wallis followed by Dunn’s test and are for data aggregated at the well level. Significance are, *** = p < 0.001, ** = p < 0.01 and * = p < 0.05. **E.** XZ MIPs from exemplar fields of view showing separation of cells into proximal and distal groups for *control* cells, and cells with depletion of *FARP1* or *TIAM2* as indicated. **F.** Stacked projections of cells with nuclei proximal or distal to the coverslip for control cells and cells depleted for *FARP1* or *TIAM2* as indicated.

To understand how specific Rho-regulators control protrusivity when positioned near or far from a rigid substrate, we compared median cell protrusivity of control and Rho-regulator depleted WM266.4 melanoma cells. To account for the effect of cell microenvironment we made these comparisons separately for proximal and distal cells (Fig 4C & D). In cells proximal to the coverslip, depletion of *FARP1*, *TIAM2*, *DOCK5* and *RND3* each decreased protrusivity (Fig 4D). In the distal context only, cells depleted for *TIAM2* had reduced protrusvity compared to the control WM266.4 melanoma cells (Fig 4D). *TIAM2*, the Rho-regulator that controlled protrusivity in distal cells, also controlled protrusivity in proximal embedded cells, suggesting control of a particular shape process near to the glass coverslip is associated with the ability to control the same process far from the glass coverslip. We noted that *RND3* and *DOCK5* depleted cells both appeared to be associated with a reduction in cell protrusivity but also cell number (Fig 4C). Compared to control, the average number of cells in wells depleted of *DOCK5* and *RND3* was reduced to approximately 65 and 77 percent, respectively. Therefore to focus on changes in protrusivity that were directly related to shape control without complications introduced by cell survival, we continued our analysis with *FARP1* and *TIAM2*.

We visualised protrusivity in *FARP1* and *TIAM2* depleted cells using CAAX-GFP signal from proximal and distal cells (Fig 4E) as well as through stacked-MIPs (Fig 4F). This confirmed that *FARP1* depleted cells are round when close to the coverslip, but have protrusivity similar to control cells when positioned away from the coverslip (Fig 4E & F). In contrast, *TIAM2* depleted cells were round in both environmental contexts (Fig 4E & F). Taken together these results suggest that in our collagen system control of protrusivity is environment-stiffness dependent for *FARP1* but independent of environment for *TIAM2*.

### Scale of regulation of protrusivity

Next we looked to find the distance over which protrusivity changes as cells are positioned away from the coverslip in untreated cells. We also looked to characterise the different ranges or distances over which *FARP1* and *TIAM2* control protrusivity. To resolve this we binned cells over two micron intervals and plotted the mean cell protrusivity for distances up to 20 microns from the coverslip (Fig 5A). We found that compared to control, *FARP1* and *TIAM2* are each required for cell protrusivity in cells with nuclei positioned up to 7-8 microns from the glass coverslip, but that for distances beyond this only *TIAM2* is required for protrusivity (Fig 5A & B).

**Fig 5.**
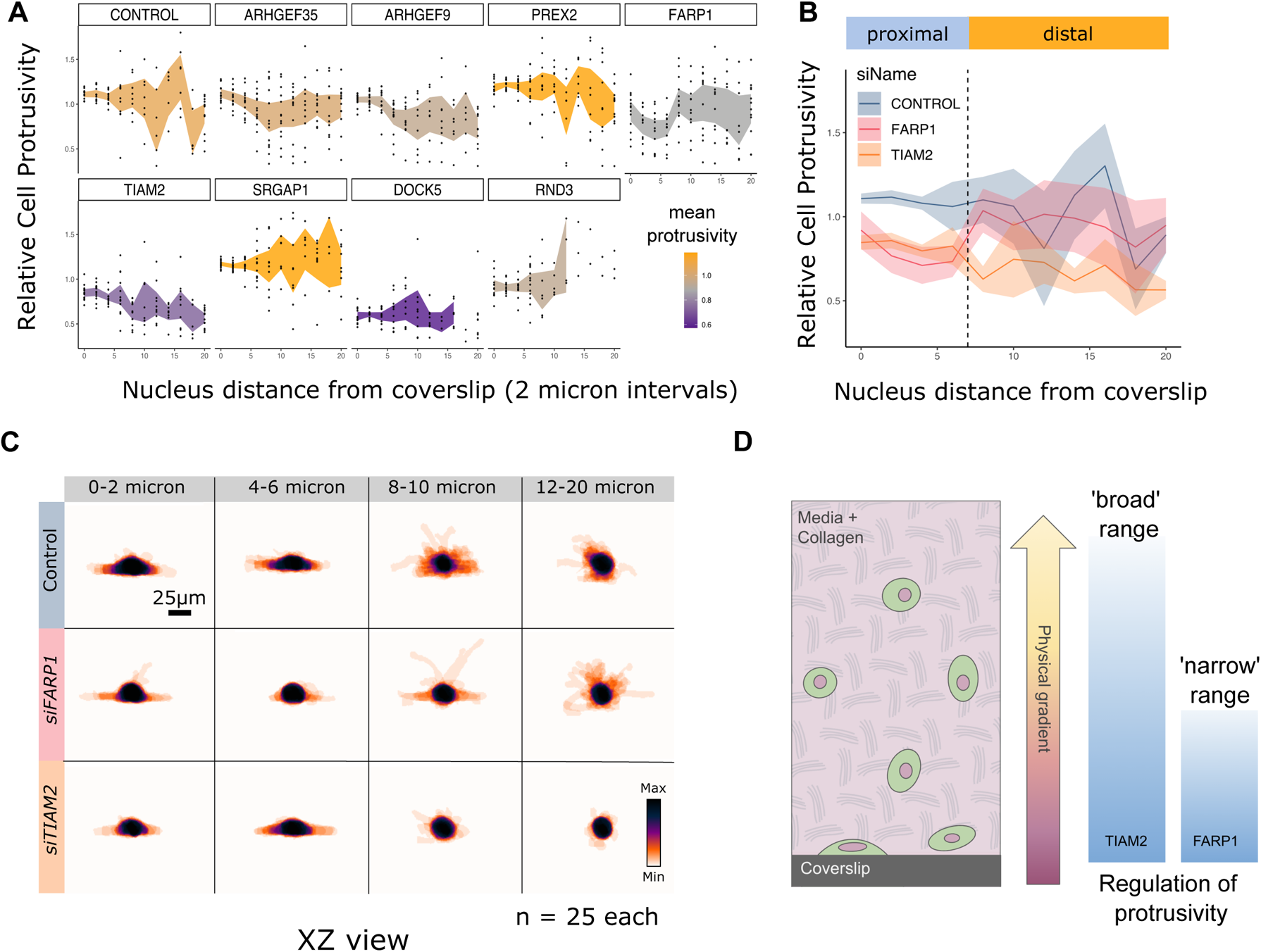
Length scale of physical regulation of protrusivity. **A.** Cell protrusivity against nucleus distance from coverslip in *control* cells, and cells depleted for Rho-regulators. Values are the mean of well level measurements. The extent of shading indicates 90 percent confidence intervals, and color of shading indicates mean protrusivity for a treatment across all distances from coverslip. **B.** Overlay of charts in A for comparison of protrusivity between *control*, and cells depleted for *TIAM2* or *FARP1*. **C.** Stacked MIPs (XZ plane) indicating changes in cell shape with distance from the coverslip. **D.** Schematic of control of protrusivity by *TIAM2* and *FARP1* over a different range of physical parameters.

To visualise the changes in protrusion that occur with distance from the coverslip, we plotted stacked-MIPs in the XZ plane at a range of intervals for cells within the first 20 microns from the coverslip (Fig 5C). This confirmed that *FARP1* depleted cells regain protrusivity beginning at distances around 8-10 microns from the coverslip, and appear similar to control cells at distances of 12 or more microns from the coverslip. Visual inspection also confirmed that *TIAM2* treated cells have a large reduction in protrusivity at all distances from the coverslip (Fig 5C).

Here our genetic perturbation data indicates that - for the gel used in this study - distances on the order of 7 microns mark a threshold, beyond which the molecular control of protrusivity is ‘handed over’ from *FARP1* to other shape regulators including *TIAM2*. This suggests a model where the control of protrusivity relies on both *TIAM2* and *FARP1* in the micro-environment close to the coverslip, but that control of protrusivity ‘switches’ to rely on *TIAM2* in the environment further away from the coverslip (Fig 5D). These context specific roles may reflect different abilities of *FARP1* and *TIAM2* to engage with distinct states of the cytoskeleton. For instance *FARP1* signalling may engage with the filamentous actin and stable integrin adhesions known to be present in coverslip cultured cells. In contrast *TIAM2* may be important in cytoskeletal states in soft gels, such as the absence of actin stress fibers.

### Control of cell height

In 2D tissue culture systems cells only contact their growth substrate at the basal surface. This means that changes in cell geometry are largely restricted to the plane of the tissue culture surface (the XY plane). In contrast, cells growing in a 3D context are embedded within their growth substrate and have greater opportunity to change geometry in the XZ plane. The axial extent, or extension of a cell and nucleus into the XZ plane (Fig 6A) have each been linked to migration and control of cell geometry [2,3,46], and the orientation of the cell and nucleus with respect to each other (Fig 6E) are connected to cell migration and disease states [46]. Due to the importance of cell and nuclear height, and cell-nuclear orientation in disease, we examined the influence of *FARP1* and *TIAM2* on these features.

**Fig 6.**
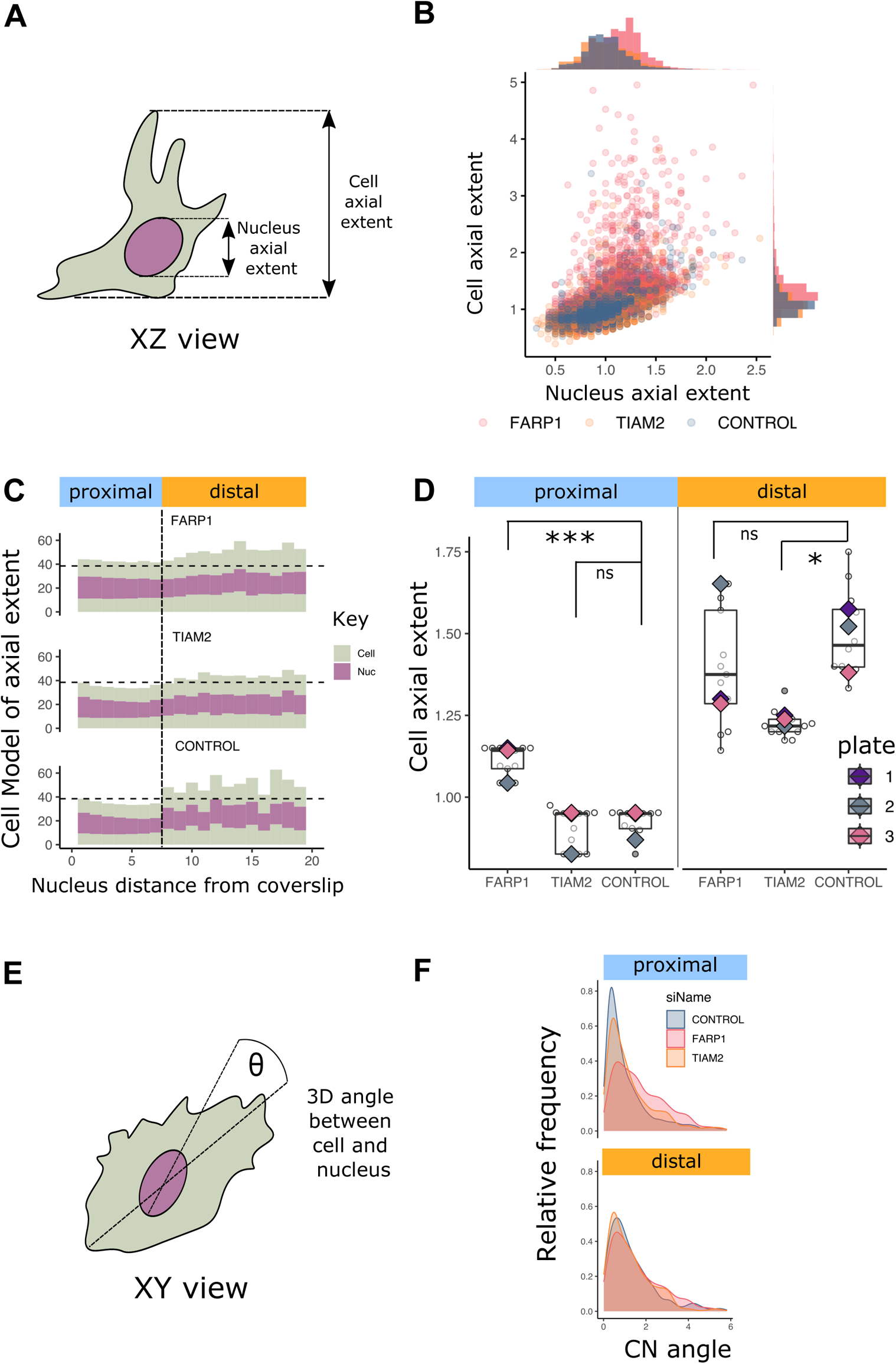
Changes in cell axial extent and cell-nuclear coupling are associated with changes in protrusivity and environment. **A.** Schematic of cell (green) and nucleus (magenta) axial extent. **B.** Plot of cell versus nucleus axial extent for cells with nuclei within 20 microns of the coverslip. Treatments are as indicated. **C.** Stacked bar-chart representation of average cell and axial extent, and nucleus position for cells at two micron intervals from the coverslip. Treatments are as indicated. **D.** Comparison of cell axial extent for the indicated genotypes. Data aggregated at the well level (boxplots and scatter-points), and plate level (rhomboids) are shown. The different colours show data from the 3 plates. Statistical tests are Kruskal-Wallis followed by Dunn’s test. **E.** Schematic of measurement for angle between cell (green) and nucleus (magenta). **F.** Estimated density plots of cell-nuclear angle in cells with nuclei proximal or distal to the coverslip. Cell treatments are indicated.

First, we plotted the relationship between cell and nucleus axial extent (the ‘height’ of the nucleus or cell) for all control, *FARP1* and *TIAM2* depleted WM266.4 melanoma cells that had their nuclei positioned within 20 microns of the coverslip (Fig 6B). This plot pooled both proximal and distal cells and indicated a positive relationship between normalised cell and nuclear height at the single cell level (Fig 6B). We also plotted frequency histograms for cell and nuclear height and noted that *FARP1* depleted cells had a shift towards increased cell and nuclear height (Fig 6B).

To visualise how cell and nuclear axial extent (height) change with distance from the coverslip we binned cells by the position of their nucleus at one micron intervals, and calculated the average cell and nucleus height (Fig 6C), as well as nuclear position. We used this information to generate ‘glyphs’ of cells that give an indication of how the relationship between cell and nucleus changes with distance of the nucleus from the coverslip (Fig 6C). This plot suggested that *FARP1* depleted cells increase their height when nuclei are within 7 microns of the coverslip, but are similar to control cells for distance beyond 7 microns. In contrast, we found that the height of *TIAM2* depleted cells are similar to control cells within the first 7 microns of the coverslip, but are reduced in height for distances beyond this. Statistical testing of the difference in height between proximal and distal cells supported these observations (Fig 6D).

The increase in height for *FARP1* depleted cells might be attributable to the concomitant increase in nuclear height (Fig 6B). Changes in nuclear height on rigid surfaces have previously been linked to maintenance of the perinuclear actin cap [46,47]. In contrast, the changes in cell height in *TIAM2* depleted cells (Fig 6C and 6D) are likely to be driven by reduced protrusivity in cells away from the coverslip (Fig 4B-D), rather than directly by changes in nuclear geometry. Considered together, these results show that *FARP1* and *TIAM2* are each required to regulate cell height, but that they regulate height in different micro-environmental contexts.

### Control of cell and nuclear coupling

Finally, we considered coupling of cell and nuclear orientation (Fig 6E and 6F). Coupling of cell and nuclear orientation is frequently observed in mammalian cell systems, where it is important for cell mechanotransduction and cell migration but this relationship breaks down in disease contexts. To measure cell and nuclear coupling we calculated the angle between the orientation of the major axis of the cell and the nucleus. We looked for changes in cell and nuclear coupling by generating probability density plots for control WM266.4 melanoma, as well as cells depleted for *FARP1* and *TIAM2*.

Consistent with previous studies, we found that in control treated WM266.4 melanoma cells there is a tight coupling of cell and nuclear orientation in proximal cells (Fig 6F). We found that this coupling is reduced in cells with their nuclei positioned distal to the coverslip (Fig 6F). Given that previous studies have also seen that changes in nuclear height are associated with breakdown of cell and nuclear coupling [47], we examined this in *FARP1* depleted cells and saw a tendency for increases in the angle between the cell and the nucleus in proximal cells.

### Comparison of cell shape in distinct environments reveals TIAM2 and FARP1 control a range of shape features but in different physical environments

Due to the importance of cell protrusivity in disease, we have focused on this metric of cell shape and found a context-dependent control of protrusvity by a subset of Rho-regulators (Fig 4A-B). However, we have found that shape differences between proximal and distal environments can be characterised by the changes in three additional shape measures (Fig 3A). Therefore we considered how each of the Rho-regulators in our study controlled these individual shape features, and whether this control is physical context specific (Fig 6A-D). To test the effect of Rho-regulators across a range of shape features, we analysed well-median values for each of our global geometry features and compared them between Rho-regulator depleted cells and control cells. Comparisons were made using Kruskal-Wallis and Dunn’s tests to compare controls to treatment.

For cells proximal to the coverslip, we found that depletion of many Rho-regulators were able to change multiple shape features. The broadest acting shape controllers in proximal cells were *FARP1*, *DOCK5*, *TIAM2* and *RND3*. The proteins that acted on fewer features were *ARHGEF9* and *ARHGEF35*. *PREX2* did not significantly change any shape features in this study (Fig 7A). In contrast, for distal cells we found that cell geometry was relatively robust to depletion of the same Rho-regulators (Fig 7B), and that fewer shape features were significantly changed by Rho-regulator depletion.

**Fig 7.**
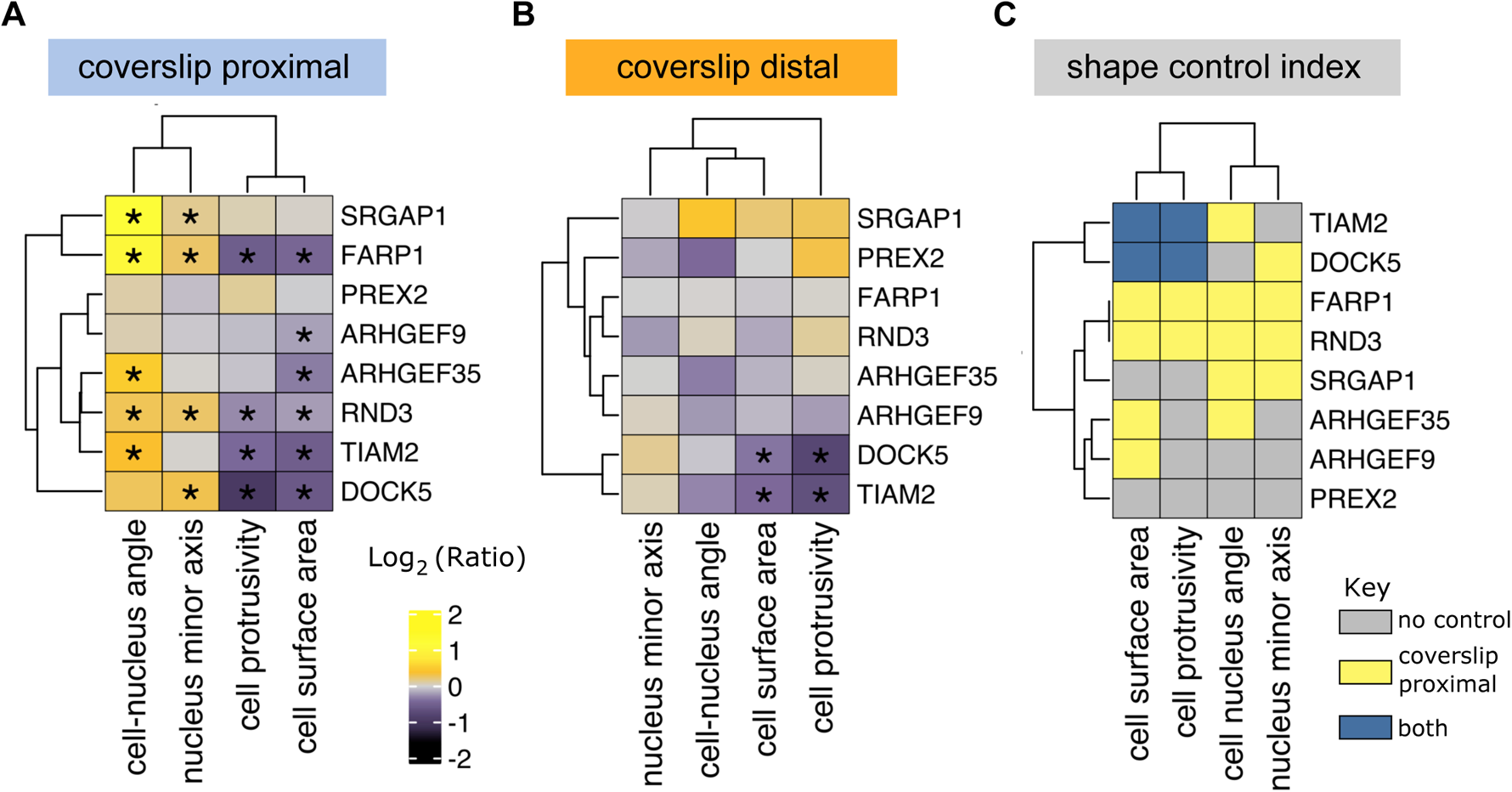
Comparison of multiple shape features across distinct environments highlights broad and specific shape controllers. **A-B.** Heatmaps indicating significant changes in cell shape features for depletion of Rho-regulators when nuclei are proximal (A) and distal (B) to the coverslip. Asterisks in A and B indicate significant change compared to control. Statistical test was Dunn’s multiple comparisons test. **C.** Logical combination of significant changes in A and B highlights genes with specific and general control of a range of shape features. For example *TIAM2* shows general shape control and affects a range of features in proximal and distal contexts. In contrast, *FARP1* shows specific shape control and influences a range of features in proximal contexts only.

To summarise the breadth and context specificity of shape control by Rho-regulators in our study, we generated a shape control matrix (Fig 7C). This highlights *DOCK5* and *TIAM2* as controlling shape features in both proximal and distal contexts (Fig 7C). In contrast, *FARP1* and *RND3* stand out as broad-acting shape regulators that significantly changed each shape feature measured, but were only effective in proximal or rigid physical regimes (Fig 7C).

## Discussion

Living cells have evolved myriad ways to change their shape in response to the physical properties of their environment. For example, a response to the environment has been linked to Rho GTPase activity and the modulation of Rho GTPases by RhoGEFs and RhoGAPs.

A full understanding of how cells change their shape in response to environmental cues is a major challenge in biology. Classic work on this problem has been conducted in 2D tissue culture paradigms, and more recently, work has been done in 3D settings. However, technical limitations have made it difficult to study the effect of RhoGEFs and RhoGAPs in multiple environments at the same time. For microscopy systems these limitations include the challenge of achieving high 3D spatial resolution and the high throughput when imaging thousands of cells simultaneously. Here, we have used ssOPM to address these challenges by imaging thousands of cells in collagen and in two distinct physical contexts. This approach creates an opportunity to understand how cells respond to different geometrical and mechanical cues, and how a genetic perturbation affects this response. We find that control treated WM266.4 melanoma cells have reduced protrusivity and become more heterogeneous when positioned away from the coverslip. A systematic depletion of Rho-Regulators revealed genes that modulate this transition. In particular our data suggest that *TIAM2* and *FARP1* ordinarily function to promote protrusions in coverslip proximal cells (Fig 4 and Fig 5). The context dependence of *FARP1* strongly suggests a role for FARP1 signalling in shape transition when cells are positioned away from rigid micro-environments.

### Stage scanning oblique plane microscopy for measuring cell shape

This assay allowed investigation of the shape of large numbers of cells at different positions in collagen. It takes advantage of many aspects of stage scanning OPM. As a light sheet technique, this method was fast. 50 volumes (corresponding to 10000 cells) were acquired in 90 minutes. This method was compatible with standard multiwell plates. This simplified the assay as standard sample preparations could be used. Nine siRNA knockdowns plus control conditions were imaged on each plate. Stage scanning allowed a large volume (4.2×0.32×0.144 mm^3^) to be imaged, corresponding to ~200 cells per volume. Notably the 144 μm z range allowed coverslip proximal and distal cells to be imaged in the same scan. This ensures values can be internally controlled. Cell segmentation and shape analysis was performed entirely in 3D. This allowed measurements of inherently 3D features including cell and nucleus axial extent.

### Validation of imaging metrics

A challenge for quantitative measurements in LSFM is that the PSF varies depending on where the sample is in the light sheet. For instance, the larger light sheet at the top and bottom of the ssOPM field of view may lead to cells appearing larger. To check for this, we imaged the same (fixed) cells positioned at different points in the light sheet by adjusting the position of the sample with respect to O1. Based on this we found that the spatially varying light sheet had a distorting effect on cell volume when imaging thin flat cells on the coverslip, and this metric was excluded from our analysis. For the other metrics used in this paper the biological effect was found to dominate over any PSF effect.

In LSFM the PSF is typically anisotropic due to the low excitation NA compared to detection. In the ssOPM system used here the resolution was (0.5×0.5×3.8 μm^3^-0.5×0.5×5 μm^3^ depending on position in the light sheet). This may lead to cells which have a dimension < 5 μm to be extended in the light sheet direction, depending on orientation. Deconvolution could be used to reduce this effect [48] but 3D deconvolution of large datasets is slow and does not take into account the spatially varying light sheet. As a simple test we eroded segmented masks with a structured element object similar to the PSF. The same cell shape changes were found with both eroded and uneroded masks. This suggests that, for this dataset, the PSF shape did not have a significant effect compared to the biological effects.

A wide range of segmentation approaches can be used in fluorescence microscopy. We tested intensity and active-contour-based segmentation. By visual inspection both methods produced good masks of cells and nuclei. We further found that we would draw the same conclusions from our data using either method.

### Cell protrusions

Cell protrusivity is essential for cell migration in normal development and metastasis during disease. However, modes of cellular protrusion are context dependent and vary with cell type, chemical and physical environment [7,49]. In 2D cell culture systems, two major modes of protrusion formation are hydrostatic blebbing and actin based protrusivity [50–52]. Hydrostatic blebbing is *RHO-ROCK* dependent and contributes to what is often described as ‘ameboid’ migration in WM266.4 melanoma cells. In contrast, actin-based protrusion is often described as ‘mesenchymal’ and relies on *RAC* regulated lamellipodia and *CDC42* regulated filopodia.

In the present study we use WM266.4 cells that have low levels of Rho GTP and produce both hydrostatic blebs and actin based protrusion but are thought to be predominately mesenchymal [7]. We also use a collagen concentration and polymerisation temperature associated with the formation of highly reticular collagen networks, and nascent, unstable integrin-based adhesions with low contractility [53].

In this setting we identified *FARP1, TIAM2, RND3* and *DOCK5* as regulating protrusivity when cell nuclei are proximal to the coverslip. In future it will be interesting to distinguish changes in the amount of hydrostatic blebbing from actin-based pseudopodial protrusions by simultaneously assessing plasma membrane and actin markers in the context of Rho-regulator depletion to determine whether either type of protrusion is specifically controlled by these Rho-regulators. Notably we also highlighted Rho-regulators where depletion increased cell protrusivity. In particular, reduction of *PREX2* and *SRGAP1* tended to increase protrusivity indicating that these Rho-regulars normally function to repress protrusivity, however these changes did not reach statistical significance in this study.

### Context specific protrusivity control

We found that most Rho-regulators were context dependent in the sense that they were more potent in controlling protrusivity close to the coverslip. This may reflect major cell biological and cytoskeletal changes that take place in response to rigidity sensing [54,55]. For example, *FARP1, RND3* and *DOCK5* may modulate protrusivity through mechanisms that depend on the abundance of integrin adhesions and filamentous actin organised into stress fibres, which are associated with rigid substrates. In contrast *TIAM2* was required for protrusion formation both proximal to and far from the coverslip and may promote protrusivity independent from environmental stiffness.

### Changes in nuclear shape and alignment

For cells in close proximity to the coverslip, the reduced protrusivity in *FARP1*-depleted cells was associated with an increase in nuclear axial extent (Fig 6C) and reduced coordination in the angle of elongation between cell and nucleus (Fig 6F). Interestingly, increased nuclear height and decreased coupling of cell-nuclear orientation were recently reported for loss of *TIAM2* [47]. In the case of *TIAM2,* increased nuclear height has been attributed to loss of nuclear capping actin, and uncoordinated cell and nuclear orientation has been attributed to dysfunction of the perinuclear actin cage [47]. In future it will be interesting to determine whether *FARP1* regulates perinuclear actin cap morphology, or is controlling nuclear axial extent by a separate mechanism.

### Effective stiffness gradients and cell and nuclear environmental sensing

Our experimental setup used cells seeded into collagen and plated on top of glass. We interpret this set-up as creating an effective ‘stiffness gradient’, where the elastic properties of the collagen proximal to the glass are influenced by the rigidity of the glass. At distances further from the glass the cells experience the greater elasticity of the collagen. Using this paradigm, we found major changes in cell shape when the base of cell nuclei are at distances on the order of 7 micrometers from the coverslip. The average nuclear diameter in our dataset was on the order of 16 micrometers. This suggests that cells respond to changes in their physical environment such as stiffness and surface geometry, over scales that are smaller than the dimensions of the nucleus.

An important question raised by our data, is whether it is the cell or nucleus that is sensing and responding to the physical environment as nuclei become positioned further away from the glass coverslip. Nuclear deformation is known to be directly linked to changes in gene expression and cell shape [56,57]. Moreover, recently new mechanisms of nuclear environment sensing have been elucidated, whereby stretching and deformation of the nuclear membrane leads to the release of calcium, increasing myosin contractility and cell shape change [2,3].

## Conclusions

In this study we use the ability of ssOPM to image thousands of melanoma cells spanning 2D and 3D collagen environments. We find cells make characteristic changes between 2D and 3D and that these changes can be modified by depletion of Rho-regulators. We find that cells in 3D environments tend to reduce their protrusivity and their protrusivity also becomes more varied and heterogeneous. Our data also suggest that cells respond to changes in environmental parameters such as stiffness and geometry, over scales that are smaller than the diameter of the nucleus. Furthermore, we identify *TIAM2* and *FARP1* as each controlling cell protrusivity but in different physical contexts. Taken together our data indicate general reliance on *TIAM2* for cell protrusivity, and a context dependent switch from *FARP1* dependent to *FARP1* independent control of protrusion between 2D and 3D settings.

## Acknowledgements

The authors wish to acknowledge the expert help of Martin Kehoe, Simon Johnson and John Murphy in the Optics Workshop of the Photonics Group of Imperial College London who contributed to the design and fabrication of components for the light-sheet microscope system.

## Abbreviations

LSFM: light sheet fluorescence microscopy
OPM: oblique plane microscopy
PSF: point spread function
ssOPM: stage scanning oblique plane microscopy

## Supplementary figure captions

**Supplementary Fig 1**

Schematic of stage scanning oblique plane microscopy (ssOPM) imaging system.

**Supplementary Fig 2**

Description of shape feature calculations. MATLAB’s regionprops3 is from the image processing toolbox.

**Supplementary Fig 3**

Examples of accepted volumes for each treatment. The same example of accepted volume for ‘Control’ treatment is also shown in Fig 1D. CAAX signal (yellow) and DRAQ5 (magenta) are shown.

**Supplementary Fig 4**

**A.** Comparison of unsupervised segmentation methods. Feature values are shown for a threshold based method and active contour method. Values were normalised to the median feature value for the plate. The trends in values are similar in both segmentation methods. Features measured using Otsu (blue) and active contour (orange) segmentation methods. **B.** Pair plots comparing features between the original (uneroded) threshold mask (blue) and an eroded version (orange). The eroded mask is chosen to take into account the anisotropic ssOPM PSF. Features are normalised to the median value for the plate.

**Supplementary Fig 5**

**A.** Measurement of the effect of the spatially varying light sheet PSF on cell shape measurements. The test plate uses shape measurements of the same cells in different axial positions with respect to the light sheet. Collagen plate data is based on the axial location of cells in collagen, within the field of view. Data points are normalised to the average cell at the bottom of the field of view across the whole dataset. A viable feature measurement should have either a normalised value close to one at all heights for the test plate, a change in the shape feature in collagen larger than on the test plate or a change in shape feature in the opposite direction to the test plate. **B.** Representative images of the test plate showing the same cells measured at different positions within the light sheet.

## References

1. Geiger B, Spatz JP, Bershadsky AD. Environmental sensing through focal adhesions. Nat Rev Mol Cell Biol. 2009;10: 21–33. doi:10.1038/nrm2593

2. Venturini V, Pezzano F, Castro FC, Häkkinen H-M, Jiménez-Delgado S, Colomer-Rosell M, et al. The nucleus measures shape changes for cellular proprioception to control dynamic cell behavior. Science. 2020;370. doi:10.1126/science.aba2644

3. Lomakin AJ, Cattin CJ, Cuvelier D, Alraies Z, Molina M, Nader GPF, et al. The nucleus acts as a ruler tailoring cell responses to spatial constraints. Science. 2020;370. doi:10.1126/science.aba2894

4. Martino F, Perestrelo AR, Vinarský V, Pagliari S, Forte G. Cellular Mechanotransduction: From Tension to Function. Front Physiol. 2018;9. doi:10.3389/fphys.2018.00824

5. Kechagia JZ, Ivaska J, Roca-Cusachs P. Integrins as biomechanical sensors of the microenvironment. Nat Rev Mol Cell Biol. 2019; 1. doi:10.1038/s41580-019-0134-2

6. Spill F, Reynolds DS, Kamm RD, Zaman MH. Impact of the physical microenvironment on tumor progression and metastasis. Curr Opin Biotechnol. 2016;40: 41–48. doi:10.1016/j.copbio.2016.02.007

7. Sahai E, Marshall CJ. Differing modes of tumour cell invasion have distinct requirements for Rho/ROCK signalling and extracellular proteolysis. Nat Cell Biol. 2003;5: 711–719. doi:10.1038/ncb1019

8. Cooper S, Sadok A, Bousgouni V, Bakal C. Apolar and polar transitions drive the conversion between amoeboid and mesenchymal shapes in melanoma cells. Mol Biol Cell. 2015;26: 4163–4170. doi:10.1091/mbc.E15-06-0382

9. Wolf K, Mazo I, Leung H, Engelke K, von Andrian UH, Deryugina EI, et al. Compensation mechanism in tumor cell migration : mesenchymal–amoeboid transition after blocking of pericellular proteolysis. J Cell Biol. 2003;160: 267–277. doi:10.1083/jcb.200209006

10. Petrie RJ, Gavara N, Chadwick RS, Yamada KM. Nonpolarized signaling reveals two distinct modes of 3D cell migration. J Cell Biol. 2012;197: 439–455. doi:10.1083/jcb.201201124

11. Sanz-Moreno V, Gadea G, Ahn J, Paterson H, Marra P, Pinner S, et al. Rac Activation and Inactivation Control Plasticity of Tumor Cell Movement. Cell. 2008;135: 510–523. doi:10.1016/j.cell.2008.09.043

12. Online Materials Information Resource-MatWeb. [cited 6 May 2021]. Available: http://www.matweb.com/index.aspx

13. Yamada KM, Sixt M. Mechanisms of 3D cell migration. Nat Rev Mol Cell Biol. 2019;20: 738–752. doi:10.1038/s41580-019-0172-9

14. Baker EL, Srivastava J, Yu D, Bonnecaze RT, Zaman MH. Cancer Cell Migration: Integrated Roles of Matrix Mechanics and Transforming Potential. PLoS ONE. 2011;6. doi:10.1371/journal.pone.0020355

15. Joshi J, Mahajan G, Kothapalli CR. Three-dimensional collagenous niche and azacytidine selectively promote time-dependent cardiomyogenesis from human bone marrow-derived MSC spheroids. Biotechnol Bioeng. 2018;115: 2013–2026. doi:https://doi.org/10.1002/bit.26714

16. McBane JE, Vulesevic B, Padavan DT, McEwan KA, Korbutt GS, Suuronen EJ. Evaluation of a Collagen-Chitosan Hydrogel for Potential Use as a Pro-Angiogenic Site for Islet Transplantation. PLOS ONE. 2013;8: e77538. doi:10.1371/journal.pone.0077538

17. Joo S, Oh S-H, Sittadjody S, Opara EC, Jackson JD, Lee SJ, et al. The effect of collagen hydrogel on 3D culture of ovarian follicles. Biomed Mater. 2016;11: 065009. doi:10.1088/1748-6041/11/6/065009

18. Tian Z, Liu W, Li G. The microstructure and stability of collagen hydrogel cross-linked by glutaraldehyde. Polym Degrad Stab. 2016;130: 264–270. doi:10.1016/j.polymdegradstab.2016.06.015

19. Jiang T, Xu G, Chen X, Huang X, Zhao J, Zheng L. Impact of Hydrogel Elasticity and Adherence on Osteosarcoma Cells and Osteoblasts. Adv Healthc Mater. 2019;8: 1801587. doi:https://doi.org/10.1002/adhm.201801587

20. Reversat A, Gaertner F, Merrin J, Stopp J, Tasciyan S, Aguilera J, et al. Cellular locomotion using environmental topography. Nature. 2020;582: 582–585. doi:10.1038/s41586-020-2283-z

21. Lämmermann T, Sixt M. Mechanical modes of ‘amoeboid’ cell migration. Curr Opin Cell Biol. 2009;21: 636–644. doi:10.1016/j.ceb.2009.05.003

22. Wolf K, Müller R, Borgmann S, Bröcker E-B, Friedl P. Amoeboid shape change and contact guidance: T-lymphocyte crawling through fibrillar collagen is independent of matrix remodeling by MMPs and other proteases. Blood. 2003;102: 3262–3269. doi:10.1182/blood-2002-12-3791

23. Brábek J, Mierke CT, Rösel D, Veselý P, Fabry B. The role of the tissue microenvironment in the regulation of cancer cell motility and invasion. Cell Commun Signal CCS. 2010;8: 22. doi:10.1186/1478-811X-8-22

24. Ridley AJ. Rho GTPase signalling in cell migration. Curr Opin Cell Biol. 2015;36: 103–112. doi:10.1016/j.ceb.2015.08.005

25. Lawson CD, Ridley AJ. Rho GTPase signaling complexes in cell migration and invasion. J Cell Biol. 2018;217: 447–457. doi:10.1083/jcb.201612069

26. Müller PM, Rademacher J, Bagshaw RD, Wortmann C, Barth C, van Unen J, et al. Systems analysis of RhoGEF and RhoGAP regulatory proteins reveals spatially organized RAC1 signalling from integrin adhesions. Nat Cell Biol. 2020;22: 498–511. doi:10.1038/s41556-020-0488-x

27. Rossman KL, Der CJ, Sondek J. GEF means go: turning on RHO GTPases with guanine nucleotide-exchange factors. Nat Rev Mol Cell Biol. 2005;6: 167–180. doi:10.1038/nrm1587

28. Kutys ML, Yamada KM. An extracellular matrix-specific GEF-GAP interaction regulates Rho GTPase crosstalk for 3D collagen migration. Nat Cell Biol. 2014;16: 909. doi:10.1038/ncb3026

29. Boureux A, Vignal E, Faure S, Fort P. Evolution of the Rho family of ras-like GTPases in eukaryotes. Mol Biol Evol. 2007;24: 203–216. doi:10.1093/molbev/msl145

30. Maioli V, Chennell G, Sparks H, Lana T, Kumar S, Carling D, et al. Time-lapse 3-D measurements of a glucose biosensor in multicellular spheroids by light sheet fluorescence microscopy in commercial 96-well plates. Sci Rep. 2016;6: 37777. doi:10.1038/srep37777

31. Dunsby C. Optically sectioned imaging by oblique plane microscopy. Opt Express. 2008;16: 20306–20316. doi:10.1364/OE.16.020306

32. Kumar S, Wilding D, Sikkel MB, Lyon AR, MacLeod KT, Dunsby C. High-speed 2D and 3D fluorescence microscopy of cardiac myocytes. Opt Express. 2011;19: 13839–13847. doi:10.1364/OE.19.013839

33. Botcherby EJ, Juskaitis R, Booth MJ, Wilson T. Aberration-free optical refocusing in high numerical aperture microscopy. Opt Lett. 2007;32: 2007–2009. doi:10.1364/OL.32.002007

34. Time-lapse 3-D measurements of a glucose biosensor in multicellular spheroids by light sheet fluorescence microscopy in commercial 96-well plates | Scientific Reports. [cited 16 Sep 2020]. Available: https://www.nature.com/articles/srep37777

35. Maioli VA. High-speed 3-D fluorescence imaging by oblique plane microscopy: multi-well plate-reader development, biological applications and image analysis. 2016 [cited 27 May 2021]. doi:10.25560/68022

36. Schindelin J, Arganda-Carreras I, Frise E, Kaynig V, Longair M, Pietzsch T, et al. Fiji: an open-source platform for biological-image analysis. Nat Methods. 2012;9: 676–682. doi:10.1038/nmeth.2019

37. Sikkel MB, Kumar S, Maioli V, Rowlands C, Gordon F, Harding SE, et al. High speed sCMOS-based oblique plane microscopy applied to the study of calcium dynamics in cardiac myocytes. J Biophotonics. 2016;9: 311–323. doi:10.1002/jbio.201500193

38. Buxboim A, Rajagopal K, Brown AEX, Discher DE. How deeply cells feel: methods for thin gels. J Phys Condens Matter Inst Phys J. 2010;22. doi:10.1088/0953-8984/22/19/194116

39. Maloney JM, Walton EB, Bruce CM, Van Vliet KJ. Influence of finite thickness and stiffness on cellular adhesion-induced deformation of compliant substrata. Phys Rev E Stat Nonlin Soft Matter Phys. 2008;78: 041923. doi:10.1103/PhysRevE.78.041923

40. Merkel R, Kirchgeßner N, Cesa CM, Hoffmann B. Cell Force Microscopy on Elastic Layers of Finite Thickness. Biophys J. 2007;93: 3314–3323. doi:10.1529/biophysj.107.111328

41. Sen S, Engler AJ, Discher DE. Matrix strains induced by cells: Computing how far cells can feel. Cell Mol Bioeng. 2009;2: 39–48. doi:10.1007/s12195-009-0052-z

42. Solon J, Levental I, Sengupta K, Georges PC, Janmey PA. Fibroblast Adaptation and Stiffness Matching to Soft Elastic Substrates. Biophys J. 2007;93: 4453–4461. doi:10.1529/biophysj.106.101386

43. Wilkinson S, Paterson HF, Marshall CJ. Cdc42-MRCK and Rho-ROCK signalling cooperate in myosin phosphorylation and cell invasion. Nat Cell Biol. 2005;7: 255–261. doi:10.1038/ncb1230

44. Yin Z, Sadok A, Sailem H, McCarthy A, Xia X, Li F, et al. A Screen for Morphological Complexity Identifies Regulators of Switch-like Transitions between Discrete Cell Shapes. Nat Cell Biol. 2013;15: 860–871. doi:10.1038/ncb2764

45. Isogai T, Dean KM, Roudot P, Shao Q, Cillay JD, Welf ES, et al. Direct Arp2/3-vinculin binding is essential for cell spreading, but only on compliant substrates and in 3D. bioRxiv. 2019; 756718. doi:10.1101/756718

46. Kim D-H, Cho S, Wirtz D. Tight coupling between nucleus and cell migration through the perinuclear actin cap. J Cell Sci. 2014;127: 2528–2541. doi:10.1242/jcs.144345

47. Woroniuk A, Porter A, White G, Newman DT, Diamantopoulou Z, Waring T, et al. STEF/TIAM2-mediated Rac1 activity at the nuclear envelope regulates the perinuclear actin cap. Nat Commun. 2018;9. doi:10.1038/s41467-018-04404-4

48. Preibisch S, Amat F, Stamataki E, Sarov M, Singer RH, Myers E, et al. Efficient Bayesian-based multiview deconvolution. Nat Methods. 2014;11: 645–648. doi:10.1038/nmeth.2929

49. Caswell PT, Zech T. Actin-Based Cell Protrusion in a 3D Matrix. Trends Cell Biol. 2018;28: 823–834. doi:10.1016/j.tcb.2018.06.003

50. Paluch EK, Raz E. The role and regulation of blebs in cell migration. Curr Opin Cell Biol. 2013;25: 582–590. doi:10.1016/j.ceb.2013.05.005

51. Petrie RJ, Yamada KM. Fibroblasts lead the way: a unified view of three-dimensional cell motility. Trends Cell Biol. 2015;25: 666–674. doi:10.1016/j.tcb.2015.07.013

52. Charras G, Paluch E. Blebs lead the way: how to migrate without lamellipodia. Nat Rev Mol Cell Biol. 2008;9: 730–736. doi:10.1038/nrm2453

53. Doyle AD, Carvajal N, Jin A, Matsumoto K, Yamada KM. Local 3D matrix microenvironment regulates cell migration through spatiotemporal dynamics of contractility-dependent adhesions. Nat Commun. 2015;6. doi:10.1038/ncomms9720

54. Nardone G, Cruz JO-DL, Vrbsky J, Martini C, Pribyl J, Skládal P, et al. YAP regulates cell mechanics by controlling focal adhesion assembly. Nat Commun. 2017;8: ncomms15321. doi:10.1038/ncomms15321

55. Gupta M, Sarangi BR, Deschamps J, Nematbakhsh Y, Callan-Jones A, Margadant F, et al. Adaptive rheology and ordering of cell cytoskeleton govern matrix rigidity sensing. Nat Commun. 2015;6: 7525. doi:10.1038/ncomms8525

56. Isermann P, Lammerding J. Nuclear Mechanics and Mechanotransduction in Health and Disease. Curr Biol CB. 2013;23. doi:10.1016/j.cub.2013.11.009

57. Dalby MJ, Riehle MO, Yarwood SJ, Wilkinson CDW, Curtis ASG. Nucleus alignment and cell signaling in fibroblasts: response to a micro-grooved topography. Exp Cell Res. 2003;284: 274–282. doi:10.1016/s0014-4827(02)00053-8

