## Supplementary figures and images for "Environmentally dependent and independent control of cell shape determination by Rho GTPase regulators in melanoma"

### Supplementary Fig 1

#### Supplementary Fig 1

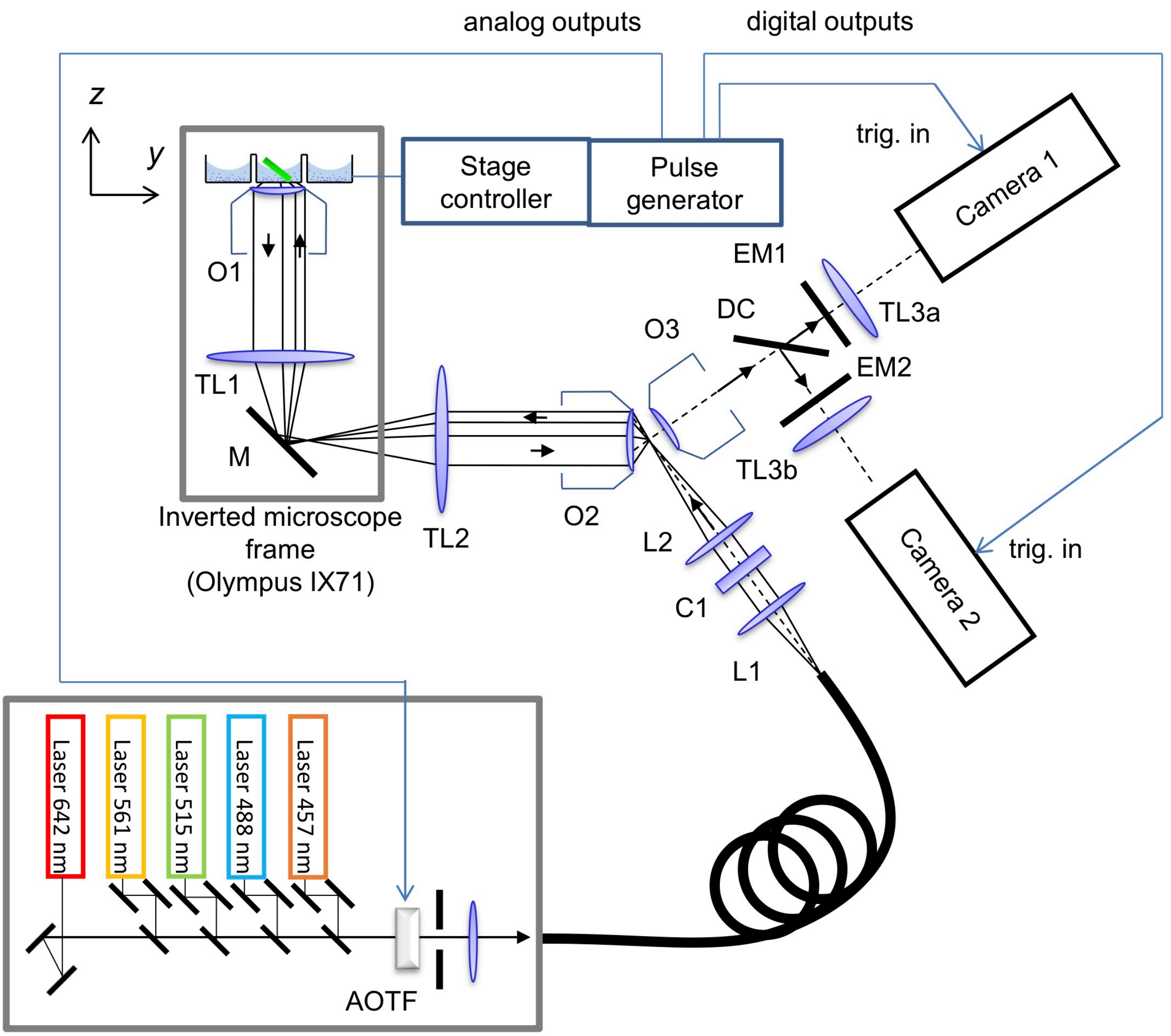

### Supplementary Fig 3

#### Supplementary Fig 3

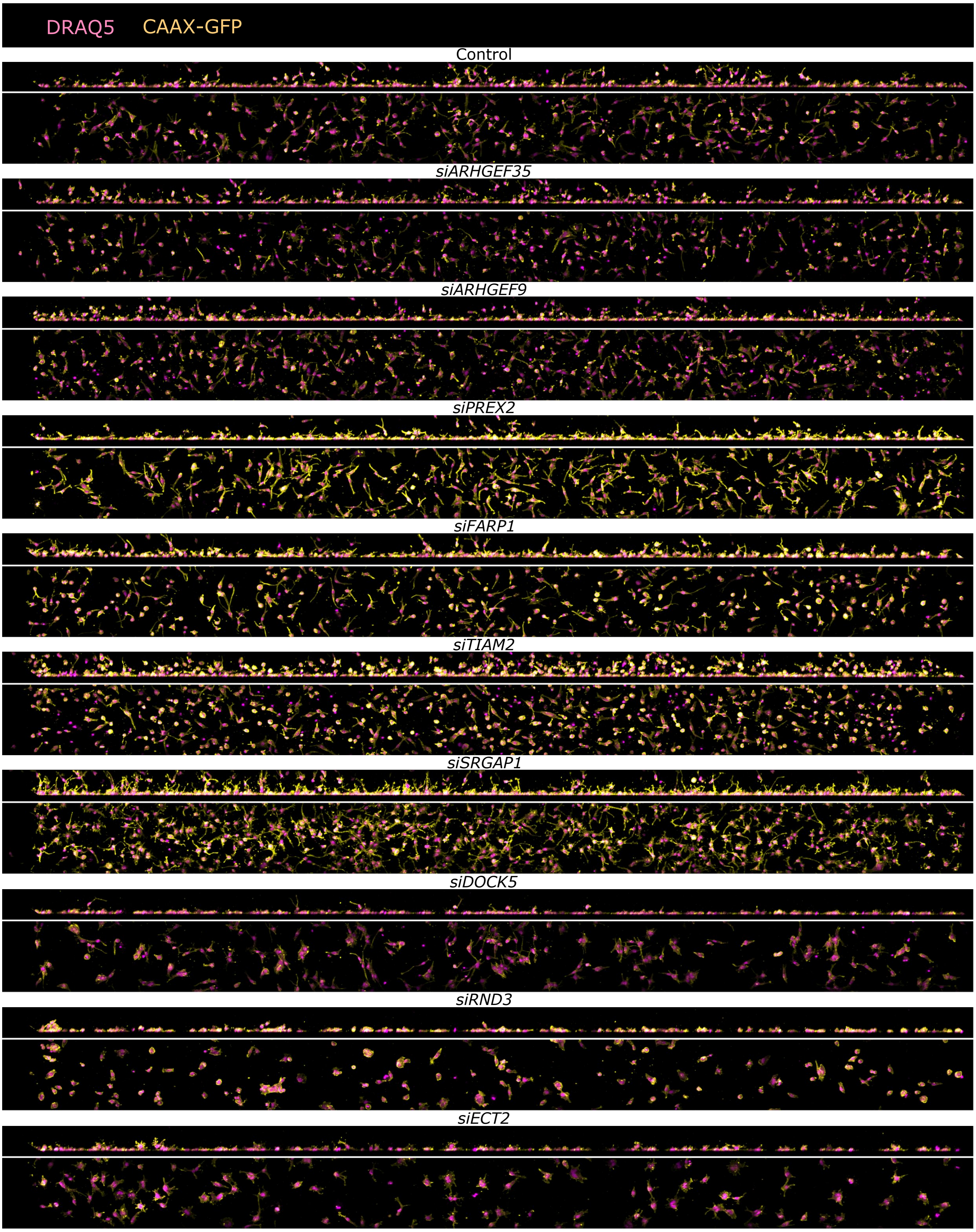

### Supplementary Fig 4

#### Supplementary Fig 4

A

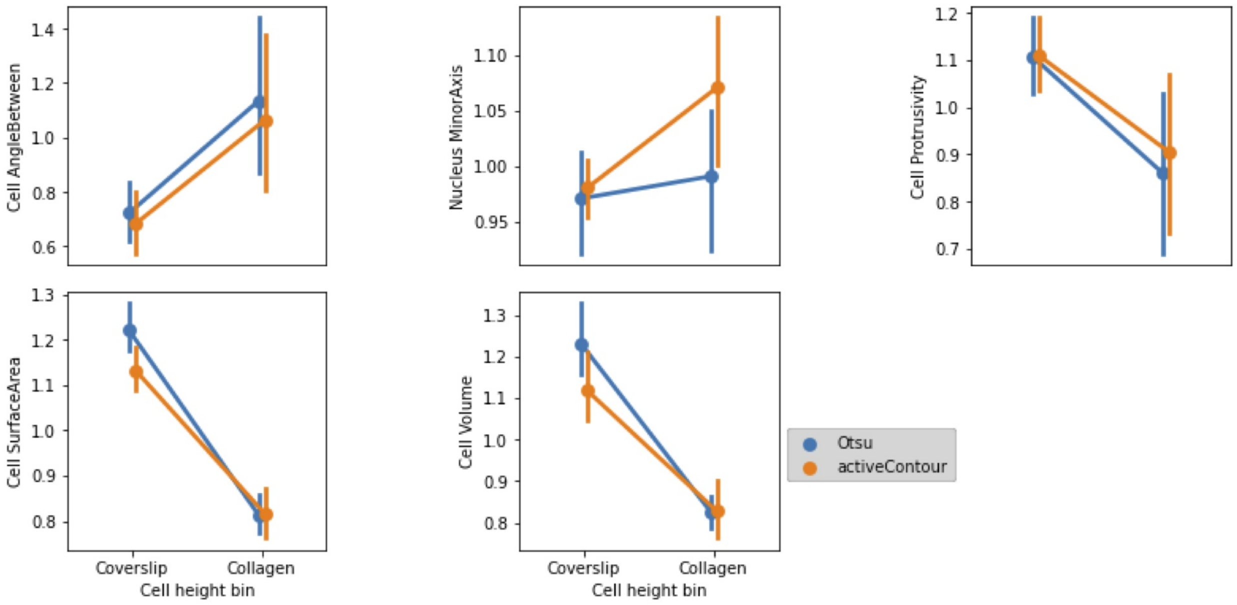


B

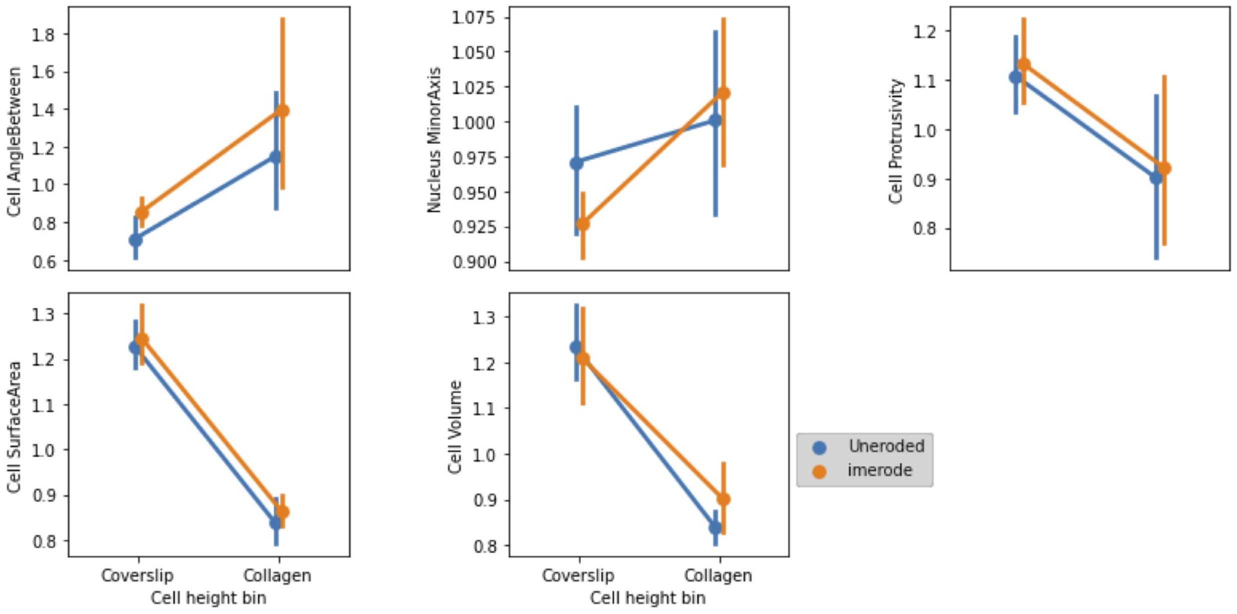

### Supplementary Fig 5

#### Supplementary Fig 5

A

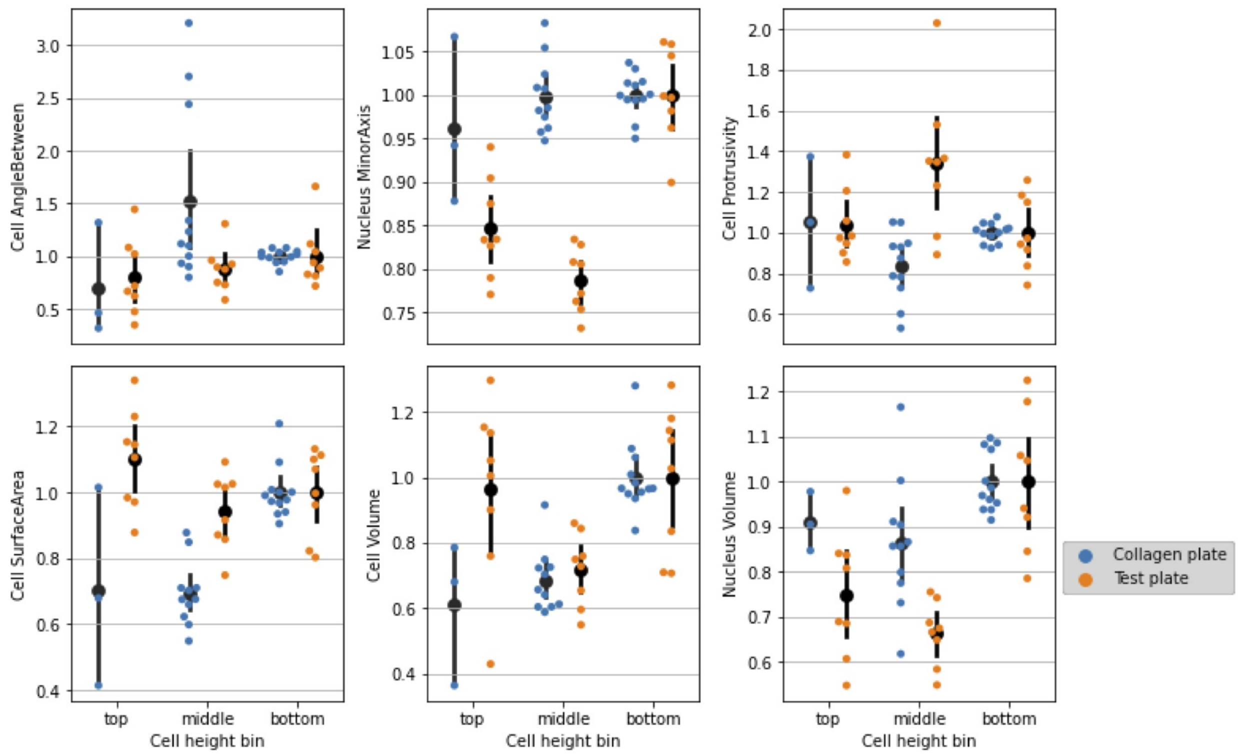


B

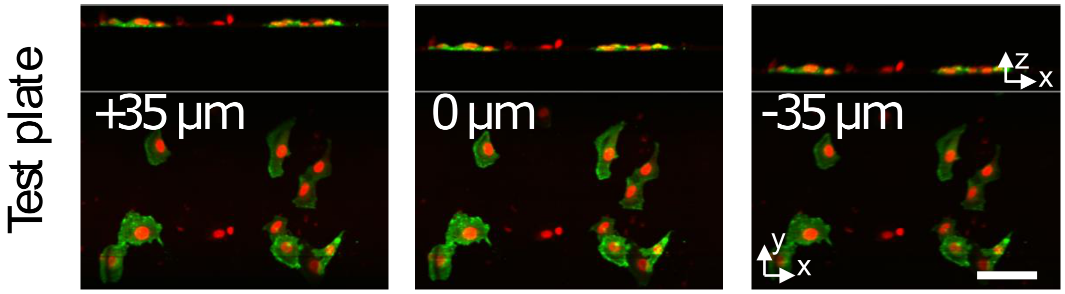
