## Supplementary Fig 2 for "Environmentally dependent and independent control of cell shape determination by Rho GTPase regulators in melanoma"

| | Name | Description | Method |
|---|---|---|---|
| Cell or nucleus metrics | Axial extent | The z range from the lowest point of the cell to the highest | From Matlab regionprops3 function. Corresponds to the height of the bounding box. |
| | Volume | Total volume of mask | Measured using Matlab regionprops3 function |
| | Protrusivity | Measure of cell spreading by comparing its volume to convex volume. | 1-volume/convex volume |
| | Surface area | Total surface area of the outside of the mask | Measured using Matlab regionprops3 function |
| | Major axis | Length of the longest ellipsoidal axis of the mask | Measured using Matlab regionprops3 function (longest PrincipalAxisLength) |
| | Minor axis | Length of the shortest ellipsoidal axis of the mask | Measured using Matlab regionprops3 function (shortest PrincipalAxisLength) |
| | Eccentricity | How circular/elliptical the plane of the minor/major axis of the mask are. 0 is a perfect circle. | $\sqrt{1-\frac{(\text{minor axis length})^2}{(\text{major axis length})^2}}$ |
| | Azimuth | Angle of the mask in the xy plane (0° = aligned along y) | "roll" from reghionprops3 orientation metric |
| | Pitch | Angle of the mask to the xy plane | "pitch" from reghionprops3 orientation metric |
| | Sphericity | How spherical the cell is (S=1, perfect sphere; Not-spherical S=0) | $\frac{\text{Surface area of a sphere with measured volume}}{\text{Measured surface area}}$ |
| | Vol2Surf | Volume to surface area ratio | $\frac{\text{Volume}}{\text{Surface area}}$ |
| | Pitch | Angle of the mask to the xy plane constrained to be between 0° and 90° | "pitch" from reghionprops3 orientation metric |
| | Second major | Length of the second longest ellipsoidal axis of the mask | Measured using Matlab regionprops3 function (middle PrincipalAxisLength) |
| | Second eccentricity | How circular/elliptical the plane of the major/second major axis of the cell are. 0 is a perfect circle. | $\sqrt{1-\frac{(\text{second major axis length})^2}{(\text{major axis length})^2}}$ |
| | Polarity | How polarised the cell is (how much longer the major axis is than the other two) 1 = highly polarised | $\sqrt{1-\frac{\text{minor axis x second major axis}}{(\text{major axis length})^2}}$ |
| | Spreading | How spread out the cell is (how much shorter the minor axis is than the other two | $\sqrt{1-\frac{(\text{minor axis length})^2}{\text{major axis x second major axis}}}$ |
| Combined metrics | Cell2nucRaio | Ratio of cell volume to nucleus volume | $\frac{\text{Nucleus volume}}{\text{Cell volume}}$ |
| | Angle between | Angle in 3D between major axes of the cell and nucleus | The orientation is converted to cartesian coordinates and then the normalised dot product is taken |
| | Orbit | Euclidean distance between cell and nucleus centre of mass | Calculated from the mask centre of mass |
| | zOrbit | Axial distance between cell and nucleus centre of mass. Positive when the cell centre of mass is above the nucleus centre of mass | Calculated from the z coordinates of the mask centre of mass |
